# The Role of Phosphoenolpyruvate Carboxylase-Protein Kinase in C_4_ Photosynthesis: Insights from *Zea mays* Mutant Analysis

**DOI:** 10.64898/2026.03.24.713513

**Authors:** Muluken Enyew, Anthony J. Studer, Russell Woodford, Maria Ermakova, Susanne von Caemmerer, Asaph B. Cousins

## Abstract

Understanding the regulation of enzyme activity involved in photosynthesis is essential for engineering enhanced carbon fixation in crops. In C_4_ plants, the enzyme phosphoenolpyruvate carboxylase (PEPC, EC 4.1.1.31) is one of the most abundant leaf enzymes and plays an essential role in photosynthetic carbon dioxide (CO_2_) fixation. The enzyme also plays a key role in central metabolism (e.g., providing intermediates to the citric acid cycle) and therefore must be highly regulated to coordinate its activity. The regulation of PEPC activity can occur allosterically by glucose 6-phosphate activation and malate inhibition, which is in part influenced by reversible phosphorylation. A specific light-dependent phosphorylation of PEPC at an N-terminal serine residue by the PEPC-protein kinase (PEPC-PK) can regulate its sensitivity to this allosteric regulation. However, the impact of this PEPC phosphorylation has not been tested in a C_4_ crop. Therefore, we created PEPC-PK mutant lines in *Zea mays* to assess the impact of PEPC phosphorylation on its allosteric regulation, photosynthesis, and growth. While the maximum PEPC activity was unchanged, PEPC in the PEPC-PK mutant plants was not phosphorylated under light and was more sensitive to malate inhibition. However, gas exchange, electron transport, and field biomass analyses showed no differences in the PEPC-PK mutant plants. These results demonstrate that in *Z. mays* PEPC phosphorylation affects enzyme sensitivity to malate *in vitro* but does not substantially alert photosynthetic performance or growth under field conditions suggesting additional regulation of PEPC activity *in planta*.

## Introduction

C_4_ photosynthesis is an efficient CO_2_ concentrating mechanism (CCM) that evolved independently in multiple plant lineages to overcome ribulose-1,5-bisphosphate carboxylase/oxygenase (Rubisco) CO_2_ limitations in C_3_ plants under conditions of high temperature, low atmospheric CO₂, and water limitation (Hibberd and Covshoff, 2010; Sage et al., 2012). By spatially separating Rubisco in bundle sheath cells and increasing CO_2_ partial pressure in that compartment, C_4_ plants reduce the rate of Rubisco oxygenation reaction and thus photorespiration (Sage, 2004; von Caemmerer, 2021). Because of this, C_4_ plants typically have high CO_2_ assimilation rates and improved nitrogen and water use efficiency (WUE) compared to C_3_ species (Ghannoum, 2009; Walker et al., 2016). These advantages of C_4_ photosynthesis have motivated global efforts to engineer C_4_ photosynthesis into C_3_ crops to enhance photosynthetic efficiency, WUE, and crop yield (Ermakova et al., 2020; Ermakova et al., 2021). However, achieving this goal requires a better understanding of C_4_ photosynthesis, including how the activity of C_4_ photosynthetic enzymes is regulated, in economically important C_4_ crops like *Zea mays* (maize) (Weber and von Caemmerer, 2010; Kajala et al., 2011).

The initial carbon fixation in C_4_ photosynthesis is performed in the cytosol of mesophyll cells by phospho*enol*pyruvate carboxylase (PEPC). Atmospheric CO_2_ is first rapidly converted by carbonic anhydrase (CA) to bicarbonate (HCO_3_^-^), which PEPC uses to catalyze the Mg^2+^ dependent carboxylation of phospho*enol*pyruvate (PEP) to form oxaloacetate (OAA). Oxaloacetate is then converted to malate or aspartate, which diffuses to the bundle sheath cells, where it is decarboxylated to release CO_2_ near Rubisco (Hatch, 1987; von Caemmerer and Furbank, 2003; Izui et al., 2004). In plants, PEPC enzymes are encoded by a small multigene family comprising two major lineages: plant type (PTPC) and bacterial type (BTPC) isoforms (O’Leary et al., 2011) Unlike PTPCs, BTPCs lack a N-terminal serine phosphorylation site, a key regulatory feature that distinguishes them from PTPCs (Sánchez and Cejudo, 2003; Carvalho et al., 2023). PTPCs typically form homotetramers and are classified as photosynthetic isoforms (C_4_ and CAM) or non-photosynthetic isoforms expressed in all C_3_, CAM, and C_4_ species (Svensson et al., 2003). Beyond carbon fixation, PEPC contributes in diverse process including osmotic adjustment, stomatal conductance, seed development, pH homeostasis, as well as central carbon and nitrogen metabolism (Britto and Kronzucker, 2005; Cousins et al., 2007; Caburatan and Park, 2021; De la Osa et al., 2022).

PEPC activity is tightly regulated by different allosteric metabolites, notably glucose-6-phosphate (G-6P) as an activator and malate as a feedback inhibitor (Doncaster and Leegood, 1987; Muñoz-Clares et al., 2020). This regulation can be further modulated by reversible phosphorylation in response to light. Specifically, in C_4_ plants, PEPC is phosphorylated in the light at the conserved N-terminal serine residue by PEPC-protein kinase (PEPC-PK) in an ATP-dependent manner (Jiao and Chollet, 1990). The signal transduction pathway leading to PEPC-PK expression is dependent on the Calvin cycle activity and involves cytosolic pH shift, inositol-1,4,5-trisphosphate production, Ca^2+^ signaling and transcriptional regulators (Jiao et al., 1991; Vidal and Chollet, 1997; Hartwell et al., 1999). The phosphorylation of PEPC by PEPC-PK reduces the enzyme’s sensitivity to malate inhibition rendering it more active *in vitro* (Vidal and Chollet, 1997). In the dark, PEPC in C_4_ plants is dephosphorylated by a specific protein phosphatase 2A (Carter et al., 1990), increasing the enzyme’s sensitivity to malate and making it less responsive to G-6P. The light-induced phosphorylation is thought to be required for maintaining activity at high concentrations of malate in the mesophyll, needed to sustain diffusion of malate to the bundle sheath cells (Bakrim et al., 1993; Vidal and Chollet, 1997; Nimmo, 2003). This fine-tune regulation of PEPC activity is likely needed to coordinate the supply of CO_2_ into the bundle sheath cells with the metabolic demand by Rubisco and energy availability, particularly under dynamic light conditions (Bakrim et al., 1993; Vidal and Chollet, 1997; Nimmo, 2003; Furumoto et al., 2007).

A limited number of studies have investigated the function of PEPC-PK-mediated phosphorylation *in planta*. Pharmacological suppression of PEPC phosphorylation using cycloheximide (inhibiting PEPC-PK synthesis) leads to a reduction in CO_2_ assimilation in *Z. mays* and sorghum whereas no comparable effect is observed in the C_3_ species wheat, where PEPC does not directly contribute to photosynthesis (Bakrim et al., 1993). In a model C_4_ dicot *Flaveria bidentis*, *PEPC-PK* knockdown and the loss of the N-terminal PEPC phosphorylation does not affect steady-state CO_2_ assimilation rates or the light induction of photosynthesis (Furumoto et al., 2007). However, *F. bidentis* is not a good model for C_4_ crops, the majority of which are grasses, and further functional analysis, including under field conditions, is needed to identify the significance of PEPC-PK in C_4_ photosynthesis.

Understanding the impact of PEPC-PK and PEPC phosphorylation on C_4_ photosynthesis is critical for reconstituting C_4_ photosynthesis in C_3_ plants, which requires not only cell-specific and high-level expression of key C_4_ enzymes but also precise post-translational and allosteric regulation (von Caemmerer et al., 2012; Ermakova et al., 2020). For instance, although *Z. mays* PEPC can be phosphorylated when expressed in rice, phosphorylation occurs in the dark, which is the opposite of *Z. mays* (Fukayama et al., 2003). Little or no contribution of *Z. mays* PEPC to CO_2_ fixation in transgenic rice plants underscores the risk of futile cycling and metabolic inefficiencies when regulatory control is not properly coordinated (Fukayama et al., 2003; Taniguchi et al., 2008; Giuliani et al., 2019; Lin et al., 2020; Ermakova et al., 2021).

We investigated the physiological role of PEPC phosphorylation in *Z. mays* using mutant lines lacking functional PEPC-PK. We examine the impact of the loss of PEPC phosphorylation on the enzyme’s *in vitro* activity and malate sensitivity. Additionally, we characterize the photosynthetic performance of mutant plants under both steady and fluctuating light as well as under field conditions. Our findings reveal that PEPC-PK mutant lines maintain photosynthesis and growth conditions comparable to wild-type plants under variety of conditions. These results suggest that the phosphorylation of PEPC in the light is not the only mechanism regulating its activity and there appears to be an additional mechanism that can counter the enzyme’s increased malate sensitivity due to the loss of phosphorylation.

## Materials and Methods

### Plant material

Mutant lines were generated using an *Activator* (*Ac*) transposon tagging approach described previously (Brutnell and Conrad, 2003). Briefly, pollen from the *r-sc∶m3/r-sc∶m3* reporter line was used to pollinate plants homozygous for the *Ac* transposable element Ac.bti00194. The Ac.bti00194 donor element was located 132 kb upstream of the *phosphoenolpyruvate carboxylase kinase1* gene (ppck1; Zm00001eb258360). From these crosses, kernels were selected that had minimal aleurone pigmentation, a hallmark of the *Ac* negative dosage effect on transposition. These 3-dose kernels indicate a duplication of the donor *Ac* element, and a potential insertion event in the target gene. A total of 741 such kernels were planted and genotyped, in pools of 10 individual plants, for an *Ac* insertion in *ppck1* using methods previously described (Studer et al., 2014). Insertion events were identified using PCR with the genic primer AJS518 [5′- CGGAAGCATTTTCTCCTGAA -3′] and the standard *Ac* primer JSR05 [5′- CGTCCCGCAAGTTAAATATGA -3′]. After identification of the *Ac* insertion events, the individual plant containing the insertion event was identified in the positive pool and Sanger sequencing was used to pinpoint the *Ac* insertion site of each mutant allele. The *ppck1-m1::Ac* allele contained an *Ac* element inserted in the 5’-3’ orientation 692 bp downstream of the start codon in an exon. The *ppck1-m2::Ac* allele contained an *Ac* element inserted in the 5’-3’ orientation 245 bp upstream of the start codon, presumably in the promoter of the gene.

### Growth conditions

At Washington State University, two independent *Z. mays PEPC-PK* mutants (*ppck1-m1* and *ppck1-m2*) and their corresponding wild type controls that were recovered from ears segregating for the PEPC-PK mutant alleles (WT M1 and WT M2) were grown in a single controlled environment growth chamber (Biochambers, GRC-36, Winnipeg, MB, Canada). Each individual plant was grown in a 8 L pot containing a mixture of Sungro professional growing mix (Bark 52%, Canadian sphagnum peat moss, perlite, dolomite lime and long-lasting wetting agent RESiLIENCE) and Turface at a 10:1 volumetric ratio. The position of pots within the growth chamber were rotated daily. Plants were grown under ambient CO_2_ conditions and 14 h photoperiod with a 2-hour light ramp, reaching a maximum light intensity of approximately 800 μmol m^-2^ s^-1^ at the canopy level. The day/night temperatures were set to 28/22°C. Plants were watered three times a week during early growth stages and every day during measurements with a nutrient solution containing Scotts-Peters Professional 10-30-20 compound (2.8 g/l), micro-chelate (6.0 mg/l), magnesium sulfate (0.6 g/l), and iron chelate 330 (1.3 g/l).

### Western blot analysis

For immunodetection of the phosphorylated PEPC, 1 cm^2^ leaf discs were collected from overnight dark-adapted or 3-hour light adapted plants and immediately frozen in liquid nitrogen. The protein was immediately extracted with glass homogenizer containing 1 ml of ice cold extraction buffer [50 mM 4-(2-hydroxyethyl)-1-piperazineethanesulfonic acid (HEPES)pH 7.8 with potassium hydroxide (KOH)], 5 mM MgCl_2_, 2 mM Ethylenediaminetetraacetic acid (EDTA), 10 mM dithiothreitol (DTT), 1% (w/v) polyvinylpolypyrrolidone (PVPP), 0.1% (v/v) Triton X-100 and 20 µl ml^-1^ of protease inhibitor cocktail (P9599, Sigma)] as described by Furumoto et al. (2007). Leaf extracts were centrifuged for 1 min at 4°C at 13,300 rpm. The supernatant was collected and diluted with equal volume of loading dye [60 mM Tris-HCl, pH 7.5, 4% (w/v) sodium dodecyl sulfate (SDS), 20% (v/v) glycerol, 1% (v/v) β-mercaptoethanol and 0.1% (w/v) bromphenol blue]. The protein was then denatured at 65 °C for 10 minutes. The protein extracts were loaded on an equal leaf-area basis and separated using SDS-PAGE Novex 8% Tris-Glycine precast gels (Invitrogen, Inc) and then transferred to a nitrocellulose membrane (Thermo Fisher Scientific Inc., 0.45 µm pore size). The membrane was blocked for 1 h at room temperature in the blocking buffer (100 mM Tris-HCl, pH 7.5, 150 mM NaCl and 1% bovine serum albumin) and then incubated overnight at 4°C with a primary antibody raised against a synthetic *Z. mays* PEPC peptide including the phosphorylated Ser-15 (Ueno et al., 2000). The goat anti-rabbit secondary antibody was visualized with 5-bromo-4-chloro-3-indolyl-phosphate/Nitro blue tetrazolium chloride solution, BCIP/NBT (Sigma-Aldrich).

### PEPC activity assay

Leaf discs (1 cm^2^) were collected and extracted as described above using an extraction buffer containing 50 mM HEPES-KOH, pH 7.8, 15 mM MgCl_2_, 1 mM EDTA, 10 mM DTT, 1% (w/v) PVPP, 0.01% (v/v) Triton X-100 and 20 µl ml^-1^ of the protease inhibitor cocktail. The PEPC activity was immediately assayed at 25 °C in 600 µl reaction volume containing 15 μl of leaf extract in the absence of malate with the assay buffer containing 100 mM HEPES-KOH, pH 7.6, 10 mM MgCl_2_, 10 mM NaHCO_3_^-^, 0.2 mM NADH, 5 mM G6-P and 7.3 units malate dehydrogenase (Furumoto et al., 2007). For the sensitivity of PEPC activity to malate, the assay buffer was supplemented with different malate concentrations (0, 1, 2, 4, 5, 6 and 7 mM). For all assays, the reaction was initiated by adding 5 mM PEP and the oxidation of NADH was monitored spectroscopically by measuring absorbance at 340 nm using Evolution 300 UV-VIS spectrophotometer (Thermo Fisher Scientific).

### Gas exchange and fluorescence measurements

The youngest fully expanded leaves of five weeks old plants were used to measure the gas exchange and fluorescence measurements using a LI-6800 portable photosynthesis system (LI-COR Biosciences, Lincoln, NE, USA) equipped with 6800-01A multiphase flash fluorometer. For CO_2_ response curves (*A*-*C*_i_ curve), the leaf chamber was set at photosynthesis photon flux density (PPFD) of 1,500 µmol m^−2^ s^−1^ (90% red/10% blue actinic light), sample CO_2_ partial pressure of 36.3 Pa, leaf temperature of 28°C, vapor pressure deficit (VPD) of 1.5 kPa, and a flow rate of 400 μmol s^-1^. Leaves were first acclimated for 30 min to these chamber conditions and then the CO_2_ partial pressure was increased stepwise from 0.5, 1.8, 3.6, 5.4, 7.3, 9.1, 13.6, 18.2, 27.2, 36.3, 54.4, 68.1, 90.7, 108.9, 136.1 Pa with 2 and 3 min as the minimum and maximum waiting times during each step, respectively. The initial slope (IS, µmol CO_2_ m^-2^ s^-1^ Pa^-1^) of each *A*-*C*_i_ curve was calculated using the first eight data points, in the linear range.

To explore transient responses of gas exchange and chlorophyll fluorescence to changes in CO_2_ and irradiance, the LI-COR leaf chamber was set at PPFD of 1,500 µmol m^-1^ s^-2^ (90% red/10% blue actinic light), a reference CO_2_ partial pressure of 36.3 Pa, leaf temperature of 28°C, VPD of 1.5 kPaand a flow rate of 400 μmol s^-1^. To examine CO_2_ responses, plants were sequentially exposed to 36.3, 18.2 and 72.6 Pa CO_2_, each for 15 minutes, and the gas exchange data was logged every 30 s. For light responses, leaves were first dark adapted for 20 min inside the LI-COR 6800 leaf chamber and the minimum (F_0_) and maximum (F_M_) fluorescence levels were recorded upon an application of a saturating pulse. Gas exchange parameters were then logged every 30 s for 10 min in the dark, 30 min at 1,500 µmol m^-2^ s^-1^, 14 min at 250 µmol m^-2^ s^-1^ and 14 min at 1500 µmol m^-2^ s^-1^. Chlorophyll fluorescence parameters (F, steady-state fluorescence; F_M_’, maximum fluorescence level under light) were recorded every 2 min upon an application of a multiphase saturating pulse of 10,000 µmol m^-2^ s^-1^. The effective quantum yield of PSII (ϕ_II_) was calculated as (F_M_’ – F) / F_M_’ (Genty et al., 1989), non-photochemical quenching (NPQ) was calculated as (F_M_ – F_M_’) / F_M_’ (Bilger and Björkman, 1990). The quantum yield of CO_2_ assimilation (ϕ_CO2_) was calculated by dividing the rate of CO_2_ assimilation (corrected for respiratory losses) by the rate of absorbed photons (Q_abs_) (Fryer et al., 1998).

### Analysis of quantum yields within photosystems under fluctuating light

For concurrent analysis of quantum yields within both photosystems, seeds were germinated in peat pots containing seed and cutting mix (Osmocote, Scotts, Bella Vista, Australia). 1 week following germination, peat pots were transplanted into 3.3 L pots filled with a 3:1 (v:v) mixture of perlite to seed and cutting mix supplemented with 5 g L^−1^ of controlled-release fertilizer (Osmocote, Scotts, Bella Vista, Australia). Plants were grown in a controlled-environment chamber with a 16 h light/8 h dark photoperiod, an irradiance of 350 µmol m^−2^ s^−1^, 28°C day, 22°C night, and 60% humidity. The youngest fully expanded leaf of 3–4-week-old plants was used for all analyses. Simultaneous measurements of Chlorophyll fluorescence and the redox state of P700 (the reaction center of PSI) were performed using DUAL-KLAS-NIR (Heinz Walz, Effeltrich, Germany) under a red actinic light using saturating pulses of 12,000 μmol m^-2^ s^-1^. Leaves were dark-adapted for 30 min before a saturating pulse was applied to obtain F_M_ and F_0_. Following this, the maximal P700^+^ signal (P_M_) was recorded upon the application of a saturating pulse at the end of an 8-second far-red light (720 nm) illumination, and the minimal P700^+^ signal (P_0_) was recorded directly after the saturating pulse in the dark. Leaves were then acclimated to 400 µmol m^−2^ s ^−1^ for 15 minutes and afterward subjected to 3 min intervals of light fluctuating between 2,000 and 400 µmol m^−2^ s ^−1^. A saturating pulse was applied every 30 s to monitor F_M_’, F, P_M_’ (the maximum level of P700^+^ under light), and P (the steady-state P700^+^ level). This allowed for tracking of the partitioning of absorbed light energy within PSII between the photochemical [ϕ_II_ = (F_M_′ − F)/F_M_′] and non-photochemical reactions, including the regulated [ϕ_NPQ_ = (F_M_ − F_M_′)/F_M_] and nonregulated (ϕ_NO_ = F/F_M_) fractions (Kramer et al., 2004). The photochemical yield of PSI [ϕ_I_ = (P_M_’ – P) / (P_M_ – P_0_)], and the non-photochemical yields of PSI due to limitation at acceptor side [ϕ_NA_ = (P_M_ – P_M_’) / (P_M_ – P_0_)] or donor side [ϕ_ND_ = (P – P_0_) / (P_M_ – P_0_)] were calculated as described in Klughammer and Schreiber (2008).

### Plant growth under fluctuating light

For fluctuating light experiment, plants were initially grown for three weeks under constant light at 800 µmol m⁻² s⁻¹. Subsequently, the plants were transferred to either a constant daylight regime (1,300 µmol m⁻² s⁻¹) or a fluctuating daylight regime (1 min at 1300 μmol m⁻² s⁻¹ followed by 4 min at 90 µmol m⁻² s⁻¹), under a 16-h light photoperiod for an additional 9 days (Schneider et al., 2019; Om et al., 2022). Light intensity was monitored using a LI-250A light meter (LI-COR Biosciences, Lincoln, NE, USA). All other growth conditions were maintained as described above. Chlorophyll fluorescence was assessed with a LI-6800 portable photosynthesis system equipped with a 6800-01A multiphase flash fluorometer (LI-COR Biosciences, Lincoln, NE, USA). Measurements of maximum quantum efficiency of PSII [F_V_/F_M_ = (F_M_ – F_0_)/F_M_] were performed after overnight dark adaptation, and NPQ and ϕ_II_ were determined following three hours of illumination. Fluorescence parameters were recorded every other day over a 10-day period. At the end of the experiment, plant height and aboveground dry biomass were measured to assess growth responses under the different light regimes.

### Field experiment

The *ppck1-m1* and *ppck1-m2* mutants and their corresponding wild type controls were grown during the summer of 2024 at the Crop Sciences Research and Education Center located in Urbana, Illinois, USA. Nitrogen fertilizer was applied prior to planting at a rate of 157 kg/hectre. A split-plot design was used, which placed mutant and controls in adjacent plots. Each plot consisted of three 3.7m rows planted with 20 kernels with 0.8m spacing between rows and 0.9m alleys between plots. Each genotype had three replicated plots, for a total of 12 plots. 56 days after planting, eight plants were harvested from the center row of each plot to measure above ground biomass. Plants were dried at 65°C for approximately two weeks prior to measuring dry weight.

### Data Analysis

All statistical analyses were conducted in R (v4.2.1), with significance set at p < 0.05. Linear mixed-effects models were fitted using the *nlme* package (Pinheiro, 2011). Estimated marginal means (EMMs) were calculated via the *emmeans* package (Lenth, 2023), and multiple comparisons were adjusted using Tukey’s method. PEPC activity across malate concentrations was analyzed using three-way ANOVA with genotype, light condition and malate concentration as fixed factors. PEPC activity in the absence of malate was analyzed by two-way ANOVA with genotype and light condition as fixed factors, including all interactions. The IC_50_ values (malate concentrations that reduces PEPC activity by 50%) was estimated for each genotype and light condition curve using a four-parameter log-logistic model; pairwise comparison of IC_50_ values were performed using EDcomp function in package *drc* (Ritz et al., 2015). The initial slope of *A*-*C*_i_ curves (µmol CO₂ m⁻² s⁻¹ µbar⁻¹), was determined for each biological replicate by fitting a linear regression to the lowest eight *C*_i_ values. Slopes were analyzed using a linear mixed-effects model with genotype as a fixed effect and replicate as a random effect. Gas exchange and Chlorophyll fluorescence under fluctuating light were analyzed using a model with time (as a factor), genotype, and their interaction as fixed effects, and replicate as a random effect to account for repeated measurements. Dry biomass data from field-grown plants were analyzed using a linear mixed-effects model with genotype as a fixed effect and block (field plot) as a random effect. All plots were produced using the ggplot2 package (Wickham, 2016), with error bars representing the standard error of the mean (±SE).

## Results

### Growth phenotype and PEPC phosphorylation

Two independent *ppck1 Ac* insertion alleles were characterized. Despite the autonomous nature of the *Ac* transposable element, the *ppck1-m1* allele represents a stable knockout of *ppck1* because of the *Ac* insertion in the exon of the gene. However, the *ppck1-m2* allele was less stable due to the 5’ insertion location, and tended to show a knockdown phenotype with the potential of reversion to wild type. To evaluate the functional role of PEPC-PK in *Z. mays*, we analyzed two independent PEPC-PK mutants alongside their respective wild type controls. The WT and PEPC-PK mutant plants exhibited no noticeable differences in growth phenotype in constant light in the environment chambers (**Figure 1**). This suggested that disruption of PEPC-PK did not produce obvious morphological or developmental defects at the vegetative stage **(Figure 1A).** Immunoblot analysis using antibody against phosphorylated PEPC (P-PEPC) revealed a strong phosphorylation signal in light-adapted WT plants, consistent with known light induced activation of PEPC via phosphorylation. In contrast, PEPC phosphorylation was completely absent in *ppck1-m1* under light conditions, confirming the loss of kinase activity. The second mutant line (*ppck1-m2*) showed variable phosphorylation patterns, suggesting partial loss of PEPC-PK function. As expected, phosphorylation was undetectable under dark conditions in all genotypes. These results confirmed that PEPC-PK was required for light dependent phosphorylation of PEPC *in planta* **(Figure 1B).**

**Figure 1.**
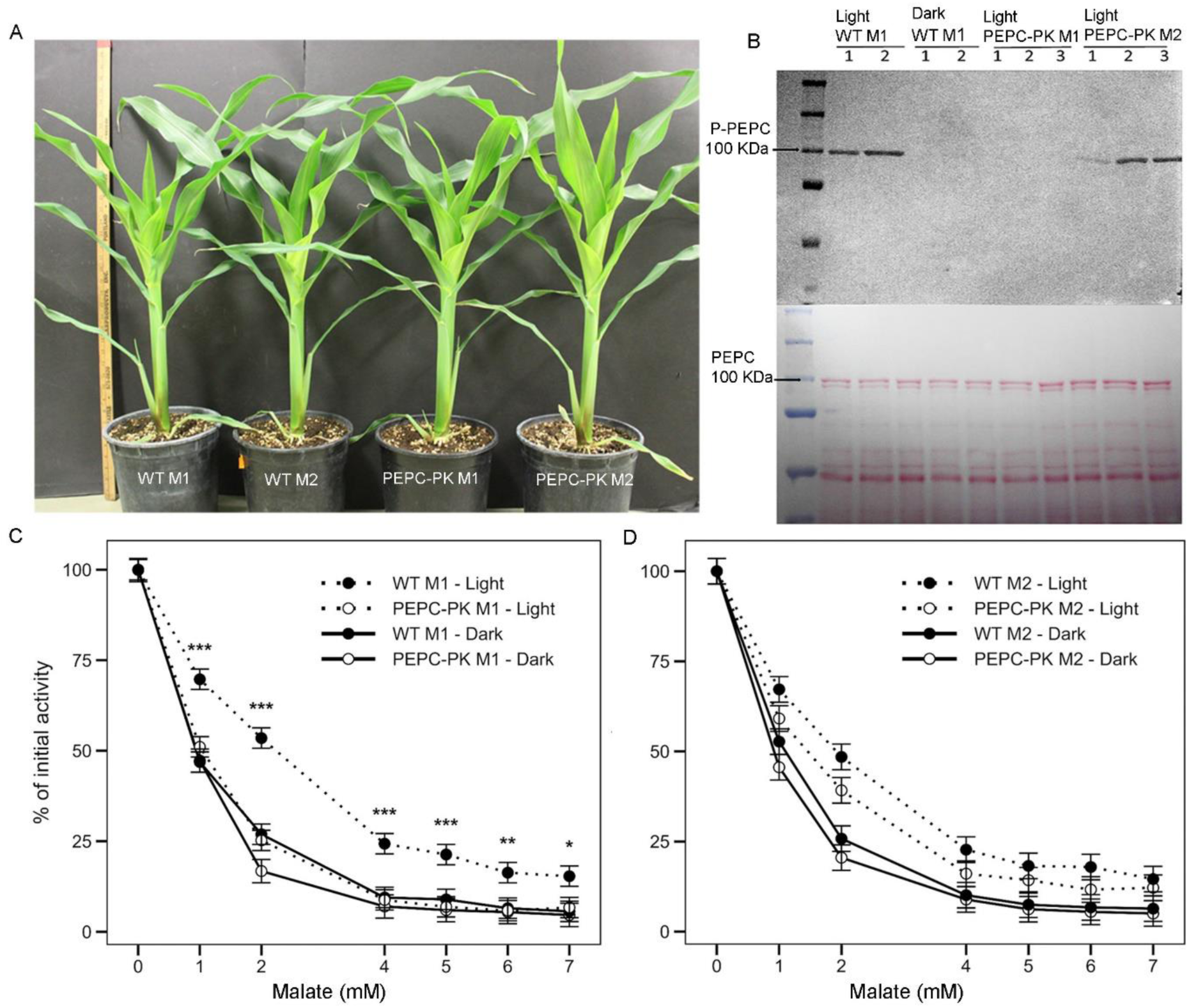
Phenotypic comparison and PEPC phosphorylation analysis in wild type and PEPC-PK mutant *Z. mays* plants. **(A)** Growth phenotype of two PEPC-PK mutants (*ppck1-m1*, *ppck1-m2*) and their corresponding wild types (WT M1, WT M2). **(B)** Immunodetection of phosphorylated PEPC (P-PEPC) in leaf protein extracts from light and dark-adapted plants. The arrow indicates the expected position of the P-PEPC band. Lanes 1, 2, 3 represents biological replicates. The lower panel shows Ponceau staining of the membrane, and the arrow points to the expected PEPC band. **(C, D)** PEPC sensitivity to malate expressed in relative units (% of activity at 0 mM malate) in WT M1 and *ppck1-m 1* (**C**), or WT M2 and *ppck1-m2*. (**D**). Mean ± SE, *n* = 4 biological replicates. Asterisks indicate significant differences between WT and *PEPC-PK* mutants under light conditions at each malate concentration (Tukey-adjusted contrasts: * *P* < 0.05, ** *P* < 0.01, *** *P* < 0.001). A three-way ANOVA revealed the significance of genotype, light condition and malate concentration and significance of two-way and/or three-way interactions among these factors, on the sensitivity of PEPC activity for malate (**Supplementary Table 1**).

### Malate sensitivity of PEPC in PEPC-PK mutants

Total PEPC enzyme activity in the absence of malate was similar between PEPC-PK mutants and their corresponding WT plants regardless of light or dark treatment (Table 1). However, across all genotypes the PEPC activity was lower in the dark treated samples than in the light adapted samples (**Table 1 and Supplementary Table 1**). In the presence of malate, PEPC activity typically decreases in the concentration-dependent manner. Activity of PEPC extracted in dark showed similar malate sensitivity between genotypes (**Figure 1C and D, and Supplementary Table 2**). The PEPC activity from light-adapted WT and *ppck1-m2* plants exhibited lower sensitivity to malate inhibition compared to dark-adapted WT plants. In contrast, malate sensitivity of PEPC extracted from the *ppck1-m1* was similar between light and dark samples (**Figure 1C and D**). The PEPC malate sensitivity expressed as IC_50_ (the inhibitor concentration of malate that reduces the PEPC activity by 50%) was not significantly different between the dark-adapted regardless of genotype (**Table 1**). The light adapted WT M1 samples showed significantly higher IC_50_ values (IC_50_ = 2.2±0.6 mM) as compared to the light adapted *ppck1-m1* samples (IC_50_ = 1.0±0.1 mM). The IC_50_ in the light adapted WT M2 (IC_50_ = 1.8±0.3 mM) and *ppck1-m2* (IC_50_ = 1.3±0.2 mM) did not differ.

**Table 1.** PEPC activity and malate IC_50_ measured in leaf extracts from light- and dark-adapted plants.

| Genotype | PEPC activity<br>($\mu\text{mol m}^{-2} \text{s}^{-1}$) | | Malate IC<sub>50</sub><br>(mM) | | Initial Slope<br>($\mu\text{mol m}^{-2} \text{s}^{-1} \text{Pa}^{-1}$) |
| --- | --- | --- | --- | --- | --- |
|  | Light | Dark | Light | Dark |  |
| **WT M1** | 296.8 ± 13.3<sup>a</sup> | 266.3 ± 14.4<sup>b</sup> | 2.2 ± 0.6<sup>a</sup> | 0.9 ± 0.1<sup>cd</sup> | 4.4±0.6<sup>a</sup> |
| ***ppck1-m1*** | 308.7 ± 14.7<sup>a</sup> | 271.0 ± 7.2<sup>b</sup> | 1.0 ± 0.1<sup>cd</sup> | 0.9 ± 0.1<sup>d</sup> | 5.0±1.0<sup>a</sup> |
| **WT M2** | 273.8 ± 11.5<sup>a</sup> | 259.4 ± 23.6<sup>b</sup> | 1.8 ± 0.3<sup>ab</sup> | 1.0 ± 0.1<sup>cd</sup> | 5.9±0.6<sup>a</sup> |
| ***ppck1-m2*** | 302.7 ± 15.5<sup>a</sup> | 264.3 ± 15.2<sup>b</sup> | 1.3 ± 0.2<sup>bc</sup> | 0.9 ± 0.1<sup>d</sup> | 3.6±0.9<sup>a</sup> |

PEPC activity was assayed at 25 °C with 5 mM PEP. Mean ± SE, $n$ = 4 biological replicates. WT M1 = wild-type 1; *ppck1-m1* = PEPC-PK mutant 1; WT M2 = wild-type 2 and *ppck1-m2* = PEPC-PK mutant 2. The malate IC<sub>50</sub> value is the inhibitor concentration of malate that reduces PEPC activity by 50%. Values are means ± SE. The initial slope of the A-C<sub>i</sub> curve ( $\mu\text{mol CO}_2 \text{m}^{-2} \text{s}^{-1} \text{Pa}^{-1}$ ) was calculated using the first eight data points from the CO<sub>2</sub> response curves of assimilation. Different superscript letters indicate significant difference among genotypes and light conditions ( $p < 0.05$ ).

### Impact of PEPC phosphorylation on gas exchange and chlorophyll fluorescence

To assess whether phosphorylation of PEPC influenced photosynthetic performance, we measured the leaf net CO₂ assimilation rate in response to intercellular CO₂ concentration (*A*-*C*_i_ curves). As shown in **Figure 2**, there were no significant differences between WT and PEPC-PK mutant lines. The initial slopes of the *A*-*C*_i_ curves, reflecting the PEPC carboxylation capacity, were estimated using the first 6-7 linear data points. The initial slopes did not differ between WT and PEPC-PK mutant plants, indicating that loss of phosphorylation did not affect PEPC activity under high light and low *C*_i_ (**Table 1**).

**Figure 2.**
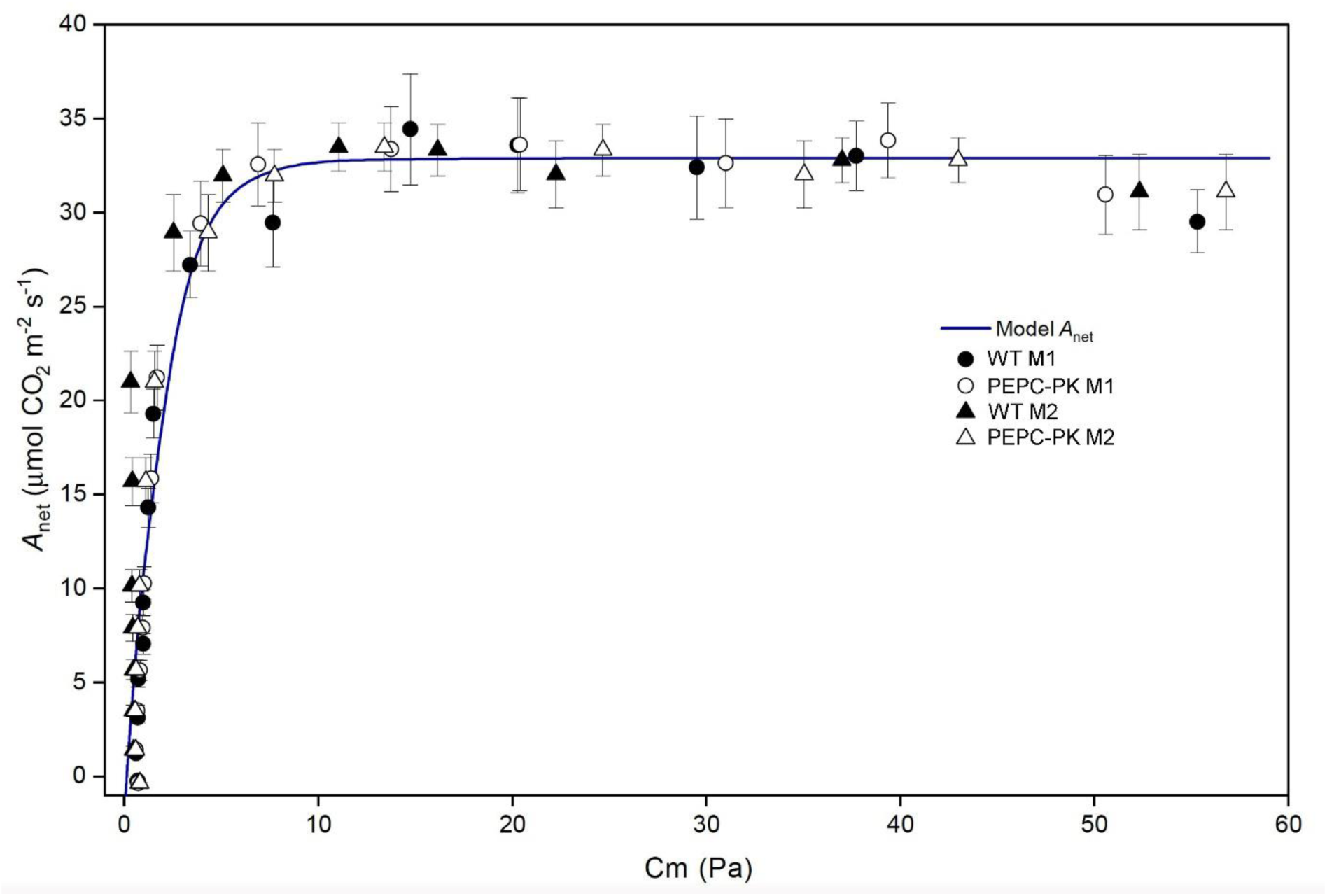
Net CO_2_ assimilation rate (A_net_) in response to the mesophyll CO_2_ partial pressure in the wild type and PEPC-PK mutant (*ppck1-m1*, *ppck1-m2*) *Z. mays* plants. Mean ± SE, *n* = 4 biological replicates. The C_4_ model curve was based on von Caemmerer (2000) using the parameter values reported by Cousins et al. (2010) for *Z. mays* with Rubisco maximum activity (*V*_cmax_) 37 μmol m^-2^ s^-1^, PEPC maximum activity (*V*_pmax_) 99 μmol m^-2^ s^-1^, bundle sheath conductance (*g*_bs_) 0.001 μmol Pa^-1^ m^-2^ s^-1^, mesophyll conductance (*g*_m_) 5 μmol Pa^-1^ m^-2^ s^-1^, and day respiration (*R*_D_) = 4 μmol m^-2^ s^-1^. No statistically significant differences were observed among genotypes (ANOVA and Tukey’s test at *P* < 0.05).

To investigate whether PEPC phosphorylation could modulate photosynthetic responses during light or CO_2_ fluctuations, we measured net CO₂ assimilation rate, stomatal conductance (*g*_sw_), quantum yield of CO₂ assimilation (ϕCO₂), effective quantum yield of Photosystem II (ϕ_II_), and non-photochemical quenching (NPQ) during transitions between different light and CO_2_ levels (**Figure 3, 4 and Supplementary Figure 1**). No differences were detected between WT and PEPC-PK mutant plants across all transitions (dark to1500 μmol m^-2^ s^-1^ to 250 μmol m^-2^ s^-1^ to 1500 μmol m^-2^ s^-1^ of light and 36.3 Pa to 18.2 Pa to 72.6 Pa CO_2_) (**Figure 3 and 4**). However, there was slightly higher *g*_sw_ in the *ppck1-m2* mutant compared to WT M2 following the transitions from 250 μmol m^-2^ s^-1^ and 1500 μmol m^-2^ s^-1^ of light and from 18.2 Pa to72.6 Pa CO_2_ (Figure 3D and Figure 4D). Responses of quantum yields of photochemical and non-photochemical reactions within photosystems to different irradiance (light curves) or to irradiance fluctuating between 400 and 2000 μmol m⁻² s⁻¹ did not differ between PEPC-PK mutant plants and corresponding WTs **(Figure 5** and **Supplementary Figure 2-4**). Collectively, these results indicated that phosphorylation of PEPC did not exert a major influence on photosynthetic performance in *Z. mays* under steady-state or fluctuating conditions.

**Figure 3.**
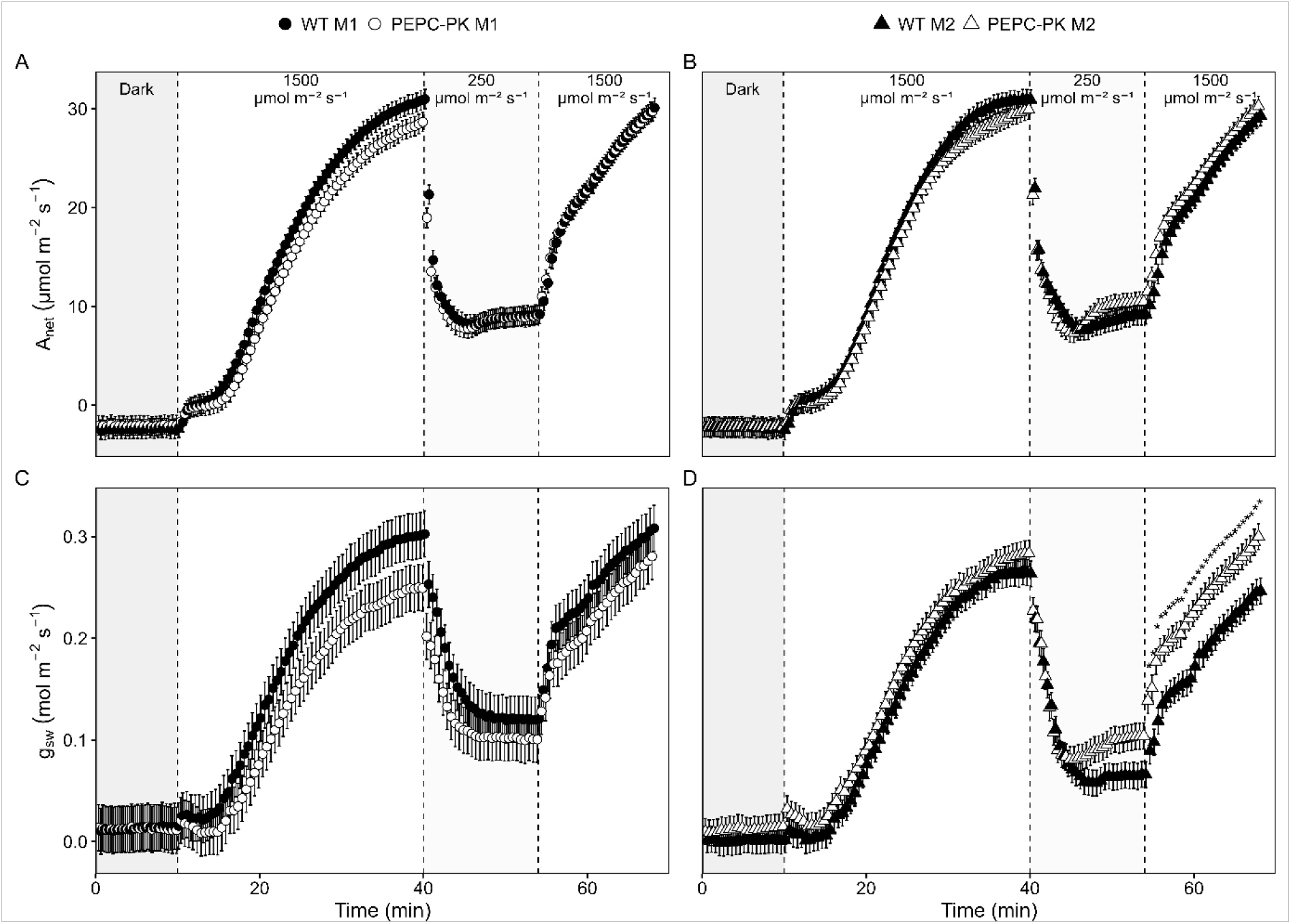
Light responses of net CO_2_ assimilation rate (A_net_) and stomatal conductance (*g*_sw_) of two *Z. mays* PEPC-PK mutants and their corresponding wild types. (**A, C**) Genotype M1: Wild type (WT M1, black circles) and mutant (*ppck1-m1*, white circles). (**B, D**) Genotype M2: Wild type (WT M2, black triangles) and mutant *ppck1-m2*, white triangles). Mean ± SE, *n* = 5-6 biological replicates. Shaded areas indicate dark periods; light intensity phases are denoted at the top of each panel. Asterisks indicate significant differences between genotypes (Tukey-adjusted contrasts at *P* < 0.05).

**Figure 4.**
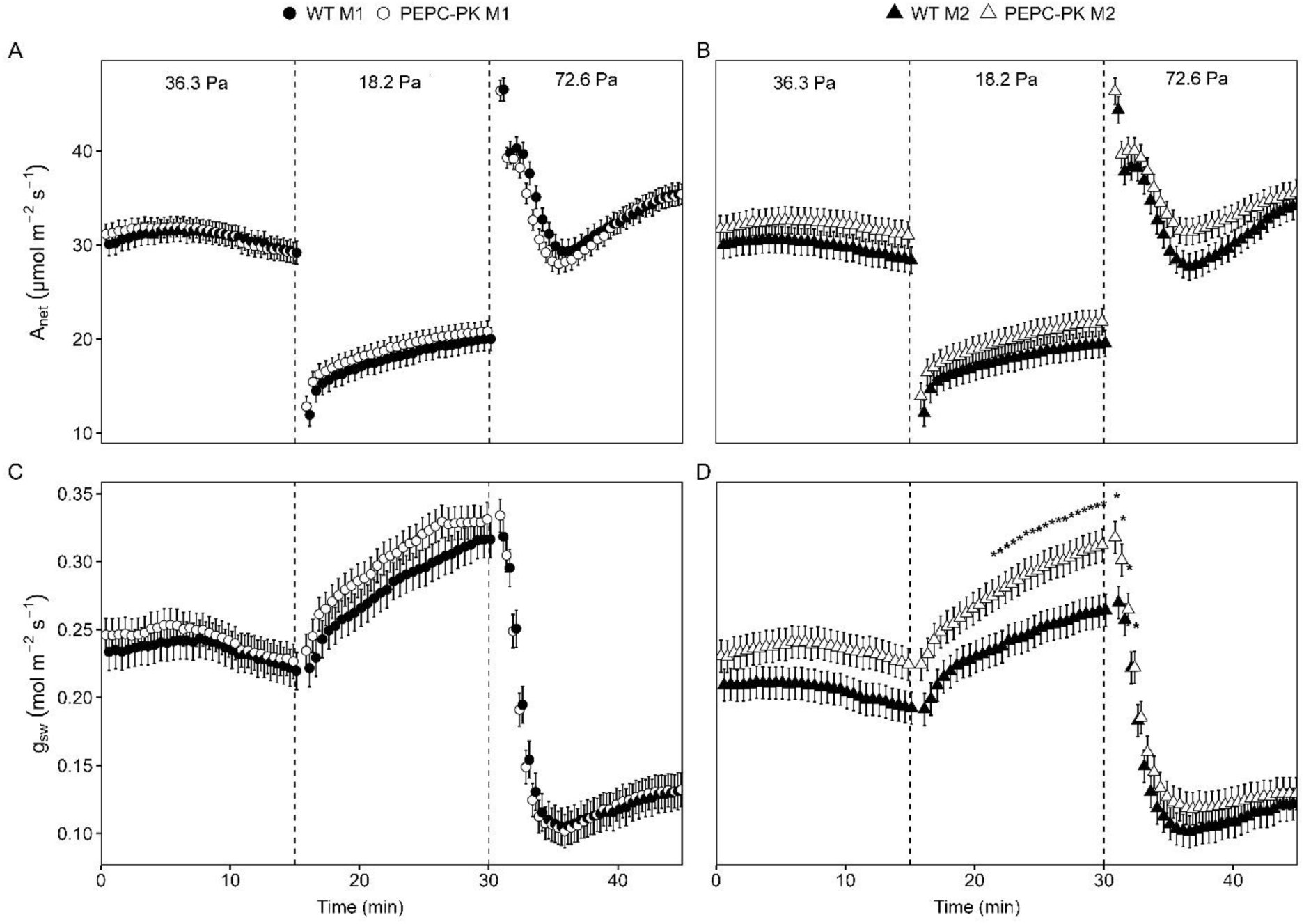
Response of net CO₂ assimilation rate (Aₙₑₜ) and stomatal conductance (*g*_sw_) to transient changes in CO_2_ of two wild type *Z. mays* genotypes and their corresponding PEPC-PK mutants. (**A, C**) Genotype M1: Wild type (WT M1, black circles) and mutant (*ppck1-m1*, white circles). (**B, D**) Genotype M2: Wild type (WT M2, black triangles) and mutant *ppck1-m2*, white triangles). Plants were exposed sequentially to 36.3, 18.2 and 72.6 Pa CO_2_, each for 15 minutes, while maintained at constant light intensity of 1500 µmol m⁻² s⁻¹. The data were collected with Portable photosynthesis systems LI-6800. Gas exchange data were recorded every 30 seconds. Mean ± SE, *n* = 5 biological replicates. Pairwise comparisons between genotypes at each time point were conducted using Tukey-adjusted contrasts. The asterisk above each points indicate significant differences at *P* < 0.05.

**Figure 5.**
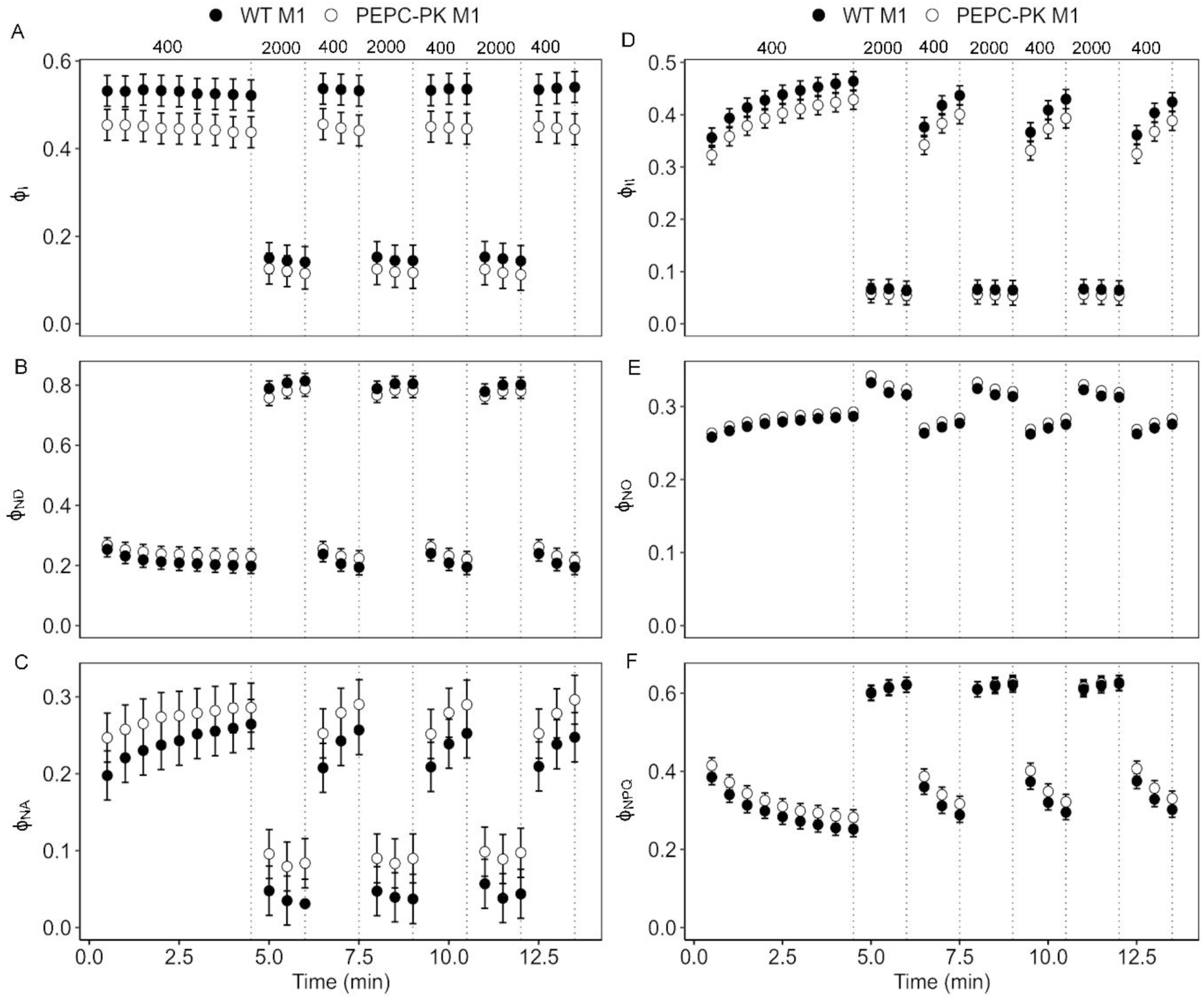
Response of Photosystem I (PSI) and Photosystem II (PSII) quantum yields in *Z. mays* wild type and PEPC-PK mutant (*ppck1-m1*) under light fluctuations. Plants were exposed to irradiance fluctuating between 400 and 2000 µmol m^-^² s^-^¹. **(A-C)** The yield of photochemical reactions in PSI (ϕ_I_), non-photochemical loss due to the oxidized primary donor of PSI (ϕ_ND_), the non-photochemical yield of PSI due to acceptor side limitation (ϕ_NA_); **(D-F)** the effective quantum yield of PSII (ϕ_II_), quantum yield of non-regulated energy dissipation (ϕ_NO_), and the yield of non-photochemical quenching in PSII (ϕ_NPQ_). Mean ± SE, *n* = 6 biological replicates. No statistical differences were found between genotypes (Tukey-adjusted contrasts at *P* < 0.05).

### Effect of fluctuating light on growth and photosynthesis

To assess the effect of fluctuating light on plant performance, WT M1 and *ppck1-m1* plants were grown under either constant or fluctuating daylight for 9 days. Growth under fluctuating daylight was slower in both genotypes compared to constant daylight but mutant and WT plants exhibited comparable growth under both light regimes, with no differences in plant height or aboveground biomass (**Figure 6A-C**). Chlorophyll fluorescence analysis revealed that the F_V_/F_M_ and ϕ_II_ declined slightly from day 1 to day 9 under both steady and fluctuating light (**Supplementary Figure 5**). In contrast, NPQ increased slightly over the same period in both genotypes and light regimes. However, no significant genotypic differences were detected for any of the fluorescence parameters under either steady or fluctuating light conditions.

**Figure 6.**
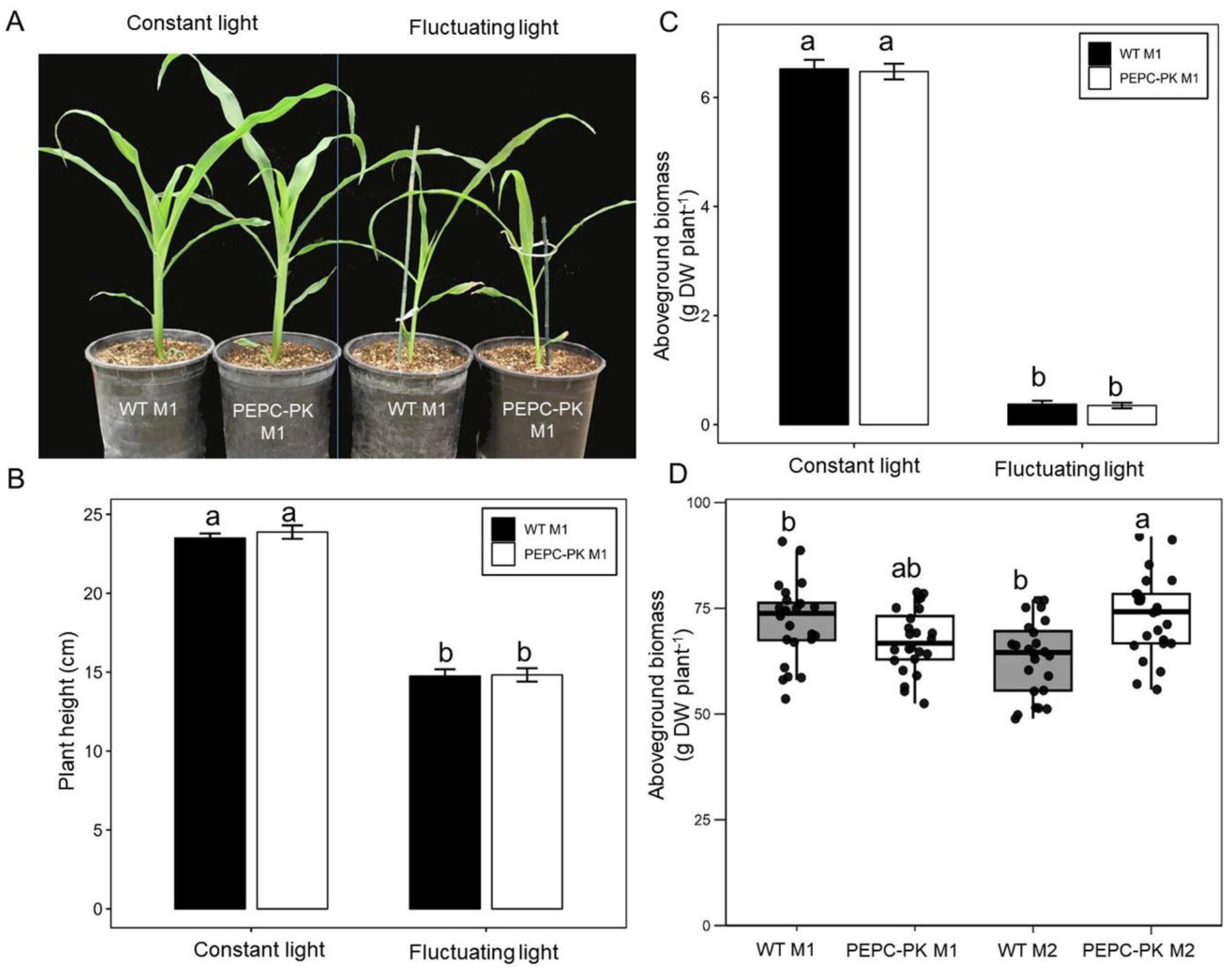
Effect of fluctuating light on growth of *Z. mays* WT and PEPC-PK mutant plants. **(A)** Growth phenotype under constant versus fluctuating daylight. **(B)** Plant height and **(C)** dry aboveground biomass. Mean ± SE, *n* = 4 biological replicates. **(D)** Aboveground biomass of field-grown *Z. mays* plants with three replicate plots per genotype. Boxes show interquartile range with the median shown by the horizontal line, whiskers denote the range and points represent individual plants. DW, dry weight. Different letters indicate significant differences between genotypes (Tukey-adjusted, *P* < 0.05).

### Field biomass and growth performance

To evaluate the impact of PEPC-PK disruption on plant growth under agronomic conditions, aboveground biomass was measured in field-grown WT and PEPC-PK mutant plants (**Figure 6D**). Across both genetic backgrounds, WT and PEPC-PK mutants produced comparable levels of biomass (**Figure 6D**). Statistical analysis revealed some variation between individual genotypes: WT M2 plants accumulated significantly less biomass than WT M1 and *ppck1-m2*, while *ppck1-m1* showed an intermediate phenotype that was not significantly different from either group. These findings indicate that loss of PEPC-PK did not impair biomass accumulation in the field.

## Discussion

Understanding how PEPC is regulated is central to modeling rates of C_4_ photosynthesis, improving photosynthetic efficiency in C_4_ crops, and introducing C_4_ traits into C_3_ crops. It has been demonstrated that PEPC activity is regulated by posttranslational modification in part through reversible phosphorylation by PEPC-protein kinase (Jiao and Chollet, 1990; Vidal and Chollet, 1997). Light-dark regulation of PEPC phosphorylation occurs in C_3_, C_4_ and CAM plants, however the temporal pattern differ among species. In C_3_ species, diurnal PEPC phosphorylation is context dependent, with studies showing variable or limited light-dark response (Fukayama et al., 2006; Meimoun et al., 2009). In CAM plants, PEPC-PK transcript abundance and PEPC phosphorylation peak at night consistent with nocturnal CO_2_ fixation (Nimmo, 2000; Boxall et al., 2017). In C_4_ leaves, light induces PEPC phosphorylation and darkness promotes dephosphorylation on the N-terminal serine (reside 15 in *Z. mays*), reflecting daytime operation of the C_4_ cycle and activation of PEPC (Terada et al., 1990; Ueno et al., 2000; Furumoto et al., 2007). This light dependent phosphorylation of this PEPC residue has been well characterized biochemically and shown to modulate the enzyme’s *in vitro* affinity for PEP, activation by G-6P, and inhibition by malate (Jiao and Chollet, 1989; Chollet et al., 1996; Nimmo et al., 2001; Nimmo, 2003). Yet despite extensive biochemical characterization, the role of PEPC phosphorylation in key C_4_ crops, particularly under fluctuating light and growth under field conditions, has not been studied.

Here, we characterized two independent PEPC-PK *Z. mays* mutants generated using *Activator* (*Ac*) insertion lines (Ac.bti00194). We assessed the impact of losing PEPC-PK on PEPC phosphorylation and the implications for *in vivo* photosynthetic performance, enzyme regulation, and plant growth in this C_4_ crop. We observed that *ppck1-m1* lacks PEPC phosphorylation at N-terminal serine in the light. This confirms the role of PEPC-PK as the kinase responsible for light-dependent phosphorylation of this residue in *Z. mays*. The phosphorylation pattern observed in the *ppck1-m2* was variable likely reflecting partial or unstable disruption of the PEPC-PK locus. As expected, phosphorylation was undetectable under dark conditions confirming that PEPC-PK is required for light dependent *in planta* phosphorylation of PEPC in *Z. mays*.

For decades, studies have shown that PEPC is highly regulated through several metabolites such as G6-P, amino acids, and malate, particularly when assayed under suboptimal pH conditions that mimic the cytosol conditions (Huber and Sugiyama, 1986; Terada et al., 1990; Duff et al., 1995). The sensitivity of PEPC to malate is regulated through the phosphorylation levels both *in vitro* or dark/light transitions (Terada et al., 1990; Jiao et al., 1991). Building on this, we examined the consequences of lack of PEPC phosphorylation at N-terminal serine in *Z. mays*. In the WT plants, the enzyme activity assays provided clear evidence that phosphorylation of PEPC reduced its sensitivity to malate inhibition under light, while the PEPC from the *ppck1-m1* mutant was more sensitive to malate inhibition. There was no difference in PEPC malate sensitivity between genotypes sampled under dark conditions, consistent with the absence of PEPC phosphorylation at night. Similarly, *PEPC-PK* knock-down in *F. bidentis* resulted in decreased PEPC phosphorylation in the light and increases the PEPC sensitivity to malate (Furumoto et al., 2007).

The high malate sensitivity of PEPC under light identified in the *ppck1-m1* mutant could be expected to negatively impact photosynthesis since the malate concentrations in the mesophyll cytosol of photosynthesizing *Z. mays* leaves can be as high as 8.4 mM (Arrivault et al., 2017; Baccolini et al., 2025). However, we found that net CO_2_ assimilation rate did not differ between the mutant and WT plants at different CO_2_ partial pressures indicating that light dependent phosphorylation of PEPC at N-terminal serine is not essential for maintaining C_4_ photosynthetic capacity *in planta* under steady-state conditions. Furthermore, the initial slope of the *A*-*C*_i_ curve, which is largely determined by PEPC activity in C_4_ plants (Peisker, 1979; Collatz et al., 1992; von Caemmerer and Furbank, 1999), was not different between genotypes. This indicated that carbon flux through PEPC at limiting and saturating CO_2_ levels was not constrained by PEPC phosphorylation status in *Z. mays*.

The tight coordination between C_4_ acid supply in mesophyll and the Calvin-Benson-Bassham demand in bundle sheath is essential for the efficiency of C_4_ pathway (von Caemmerer and Furbank, 1999), light transitions disrupt this coordination to some degree (Sun et al., 2014; Slattery et al., 2018; Arrivault et al., 2025). For instance, during high to low light transition, accumulated at high light malate or aspartate can continue to diffuse to the bundle sheath cells for decarboxylation while ATP supply drops, transiently over-pumping CO_2_ and increase bundle sheath leakiness, which cost energy for the leaf (von Caemmerer and Furbank, 1999; Slattery et al., 2018). The leakage of CO_2_ from the bundle sheath under low light transition has been reported in C_4_ plants (Cousins et al., 2006; Pengelly et al., 2010). During low to high light transitions, rapid C_4_ flux requires a build-up of large C_4_ acid pools, otherwise suboptimal CO_2_ near Rubisco elevates photorespiration(von Caemmerer and Furbank, 2003; Slattery et al., 2018). Phosphorylation of PEPC could help fine tune the activity of the CCM and supplying the needed flux of organic acids to the BSC during these transitions. However, we detected no significant differences between WT and *ppck1-m1* mutant under light transients.

We also tested whether lack of PEPC phosphorylation influences photosynthesis and growth when plants are grown under fluctuating light. In the control chambers, the fluctuating light reduced plant growth relative to steady light, but plant height, biomass and chlorophyll fluorescence did not differ between genotypes. Similarly, above ground biomass did not differ between genotypes in field grown plants. These data demonstrate that despite changes in PEPC’s phosphorylation state at the N-terminal serine and the enzyme’s increased sensitivity to malate this does not appear to impact growth or photosynthetic parameters under steady-state and fluctuating light or growth under field conditions.

The regulatory pairing of PEPC-PK and PEPC appears to have been under strong evolutionary selection during the emergency of C_4_ photosynthesis (Aldous et al., 2014), suggesting that phosphorylation serves to fine-tune PEPC activity across variable growth conditions and may facilitate specific protein-protein interactions (O’Leary et al., 2011; Grieco et al., 2012; Balparda et al., 2023). However, our findings are consistent with the earlier work in *F. bidentis*, where *PEPC-PK* knock-down reduced PEPC phosphorylation in the light but this had no significant impact on steady-state CO_2_ assimilation (Furumoto et al., 2007). In contrast, in CAM plant *Kalanchoë fedtschenkoi*, circadian-regulated phosphorylation of PEPC in the dark enhances and prolongs nocturnal CO_2_ fixation and malate accumulation (Boxall et al., 2017). These different effects of the lack of PEPC phosphorylation between C_4_ and CAM plants are likely underpinned by the central role PEPC-PK plays in circadian control of carbon metabolism in CAM plants. Additionally, CAM and C_4_ PEPCs exhibit divergent responses to metabolic effectors. While, glycine, serine, and alanine strongly activate *Z. mays* PEPC, they have negligible effect on the CAM PEPC (Nishikido and Takanashi, 1973; Bandarian et al., 1992). Critically, high concentration of glycine/alanine can fully override malate inhibition of *Z. mays* PEPC while G-6P only partially (Gao and Woo, 1996; Tovar-Méndez et al., 2000). This suggests that in C_4_ leaves, amino acid abundance can potentially neutralize malate inhibition independently of phosphorylation. Conversely, CAM PEPC mainly relies on G-6P regulation, and phosphorylation is essential for overcoming malate inhibition. Alternatively, other post translational modifications might modulate C_4_ PEPC activity *in planta* in the absence of phosphorylation. Future investigations utilizing phosphoproteomics and site-directed mutagenesis to determine if other PEPC residues can modulate PEPC activity, malate sensitivity, and photosynthetic performance.

## Conclusion

In *Z. mays*, loss of function in PEPC-PK eliminated the light dependent phosphorylation of PEPC yet total leaf PEPC activity was unchanged. Although the lack of phosphorylation increased PEPC’s malate sensitivity *in vitro*, there was no measurable effect on leaf gas-exchange, chlorophyll fluorescence, or dry biomass when plants were grown under field conditions. Overall, these results demonstrate that the light dependent phosphorylation of PEPC is not a major determinant of C_4_ photosynthetic performance in *Z. mays* suggesting there are other redundant unknown mechanisms that regulate PEPC activity in planta that may exclusively function in C_4_ plants.

## Acknowledgements

We thank Soumi Bala for having grown these mutants through quarantine in Canberra. This research was supported in part by the Australian Research Council Centre of Excellence for Translational Photosynthesis (CE140100015) to SvC; the United States Department of Agriculture—Hatch, a United States Department of Agriculture—Agriculture and Food Research Initiative grant (2019-67013- 29195) to AJS and ABC; the Office of Biological and Environmental Research in the DOE Office of Science (DE-SC0008769) to ABC; and the Australian Research Council Discovery Project (DP230100175) to MEr.

## Author contributions

ABC, MEn, SvC, AJS and MEr designed the research. MEn, RW, and AJS performed the experiments. AJS generated the mutant plants, ABC, MEn, SvC, AJS and MEr analyzed the data. MEn wrote the manuscript with constructive inputs from all authors. All authors reviewed and approved the final manuscript.

---

**Supplementary Figure 1.**
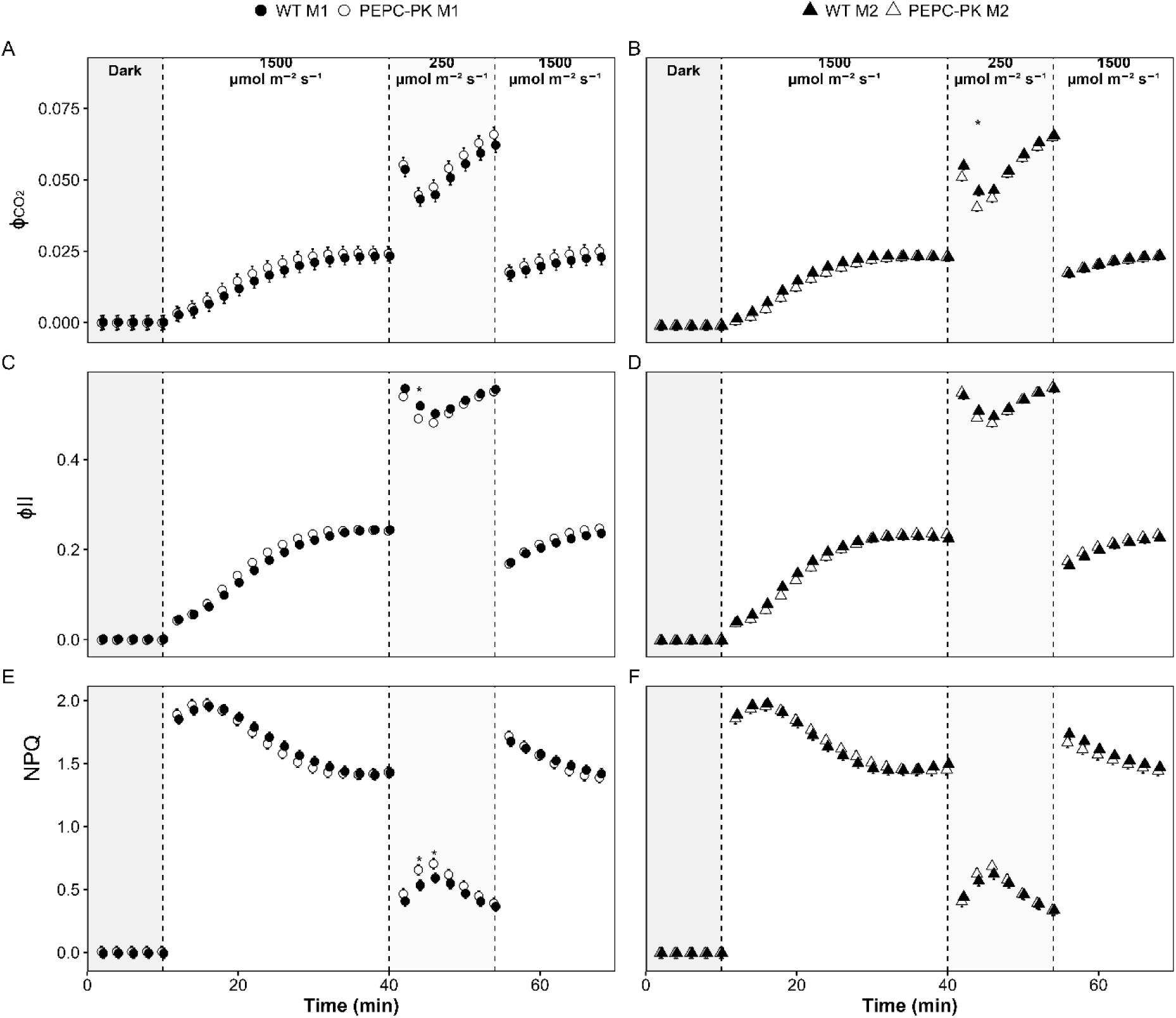
Dark-light transition of (**A**) the effective quantum yield of CO_2_ assimilation (ϕ_CO2_), (**B**) quantum yield of PSII (ϕ_II_) and (**C**) non-photochemical quenching (NPQ) for wild type (WT M1and WT M2), PEPC-PK mutant (*ppck1-m1*and *ppck1-m2*) *Z. mays* genotypes. The data were logged every 2 mins under light condition of a PPFD of 0 µmol m^-^² s^-^¹ for 10 mins,, 1500 µmol m^-^² s^-^¹ for 30 mins and 250 µmol m^-^² s^-^¹14 mins and 1500 µmol m^-^² s^-^¹ 14 mins. Points represent averages of five to six individuals ± SE. Grey boxes in each figure indicate the initial period of darkness. The grey (Dark) and white boxes (1500, 250, 1500 µmol m^-^² s^-^¹) indicate the light intensity. The asterisk above each points indicate significant differences at Tukey-adjusted *P* < 0.05.

**Supplementary Figure 2.**
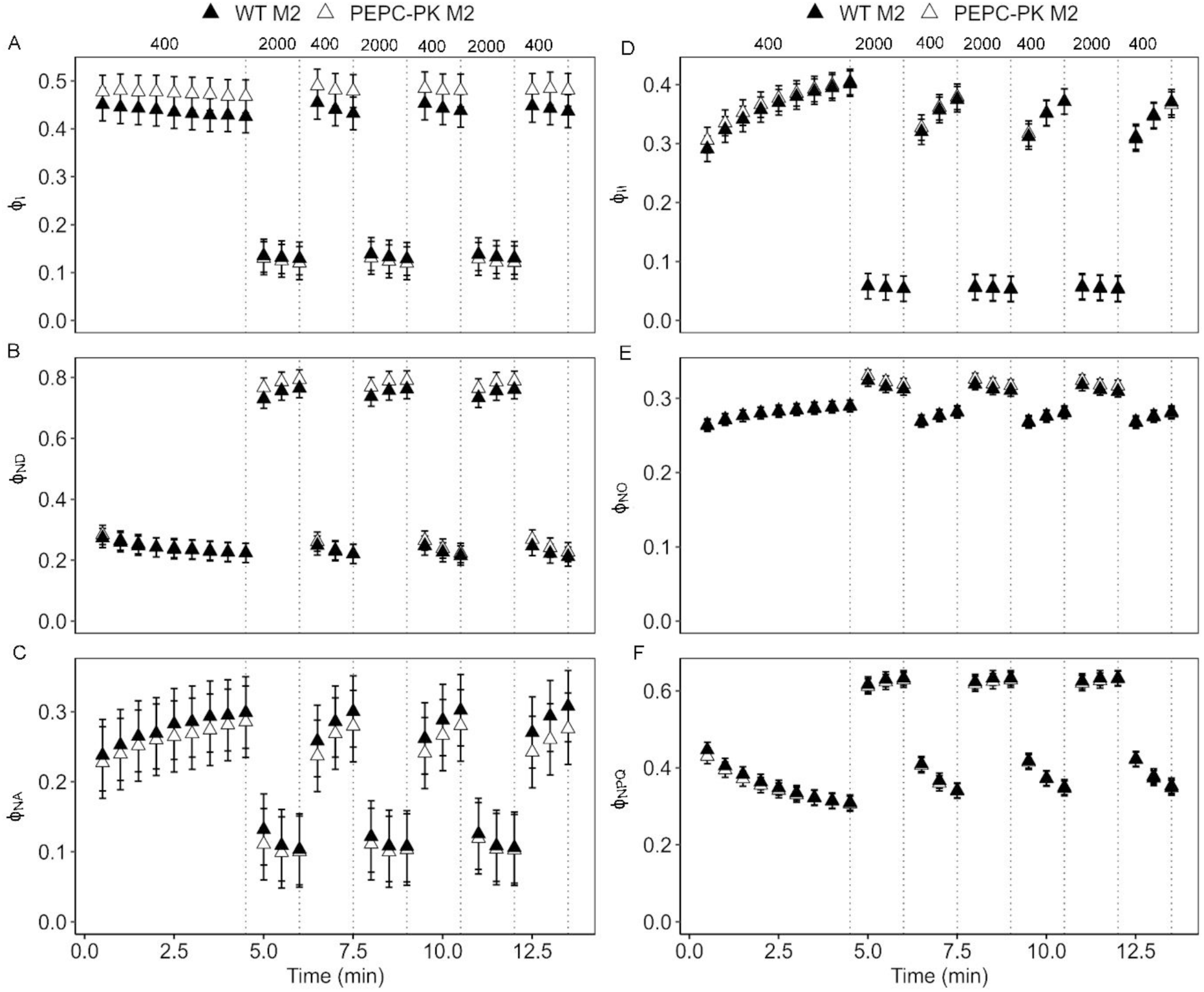
Response of Photosystem I (PSI) and Photosystem II (PSII) quantum yields in *Z. mays* wild type and PEPC-PK M2 (*ppck1-m2*) mutant under light fluctuations. Plants were exposed to irradiance fluctuating between 400 and 2000 µmol m^-^² s^-^¹. **(A-C)** The yield of photochemical reactions in PSI (ϕ_I_), non-photochemical loss due to the oxidized primary donor of PSI (ϕ_ND_), the non-photochemical yield of PSI due to acceptor side limitation (ϕ_NA_); **(D-F)** the effective quantum yield of PSII (ϕ_II_), quantum yield of non-regulated energy dissipation (ϕ_NO_), and the yield of non-photochemical quenching in PSII (ϕ_NPQ_). Mean ± SE, *n* = 6 biological replicates. No statistical differences were found between genotypes (Tukey-adjusted contrasts at *P* < 0.05).

**Supplementary Figure 3.**
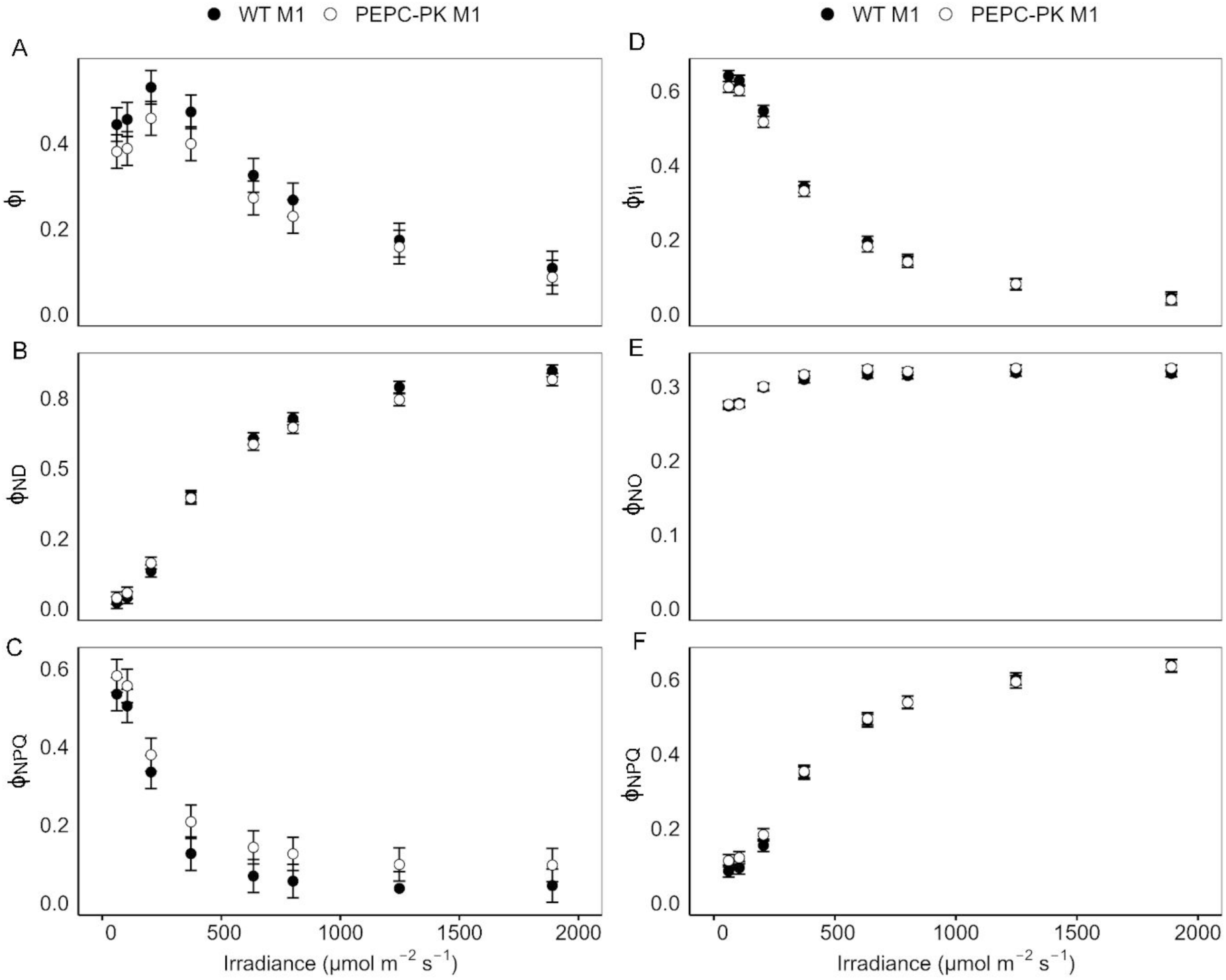
Light response curve of Photosystem I (PSI) and Photosystem II (PSII) related parameters in wild type and PEPC-PK mutant *Z. mays* genotype 1 (*ppck1-m1*). **(A-C)** The yield of photochemical reactions in PSI (ϕ_I_), non-photochemical loss due to the oxidized primary donor of PSI (ϕ_ND_), the non-photochemical yield of PSI due to acceptor side limitation (ϕ_NA_); **(D-F)** the effective quantum yield of PSII (ϕ_II_), quantum yield of non-regulated energy dissipation (ϕ_NO_), and the yield of non-photochemical quenching in PSII (ϕ_NPQ_). Mean ± SE, *n* = 6 biological replicates. No statistical differences were found between genotypes (Tukey-adjusted contrasts at *P* < 0.05).

**Supplementary Figure 4.**
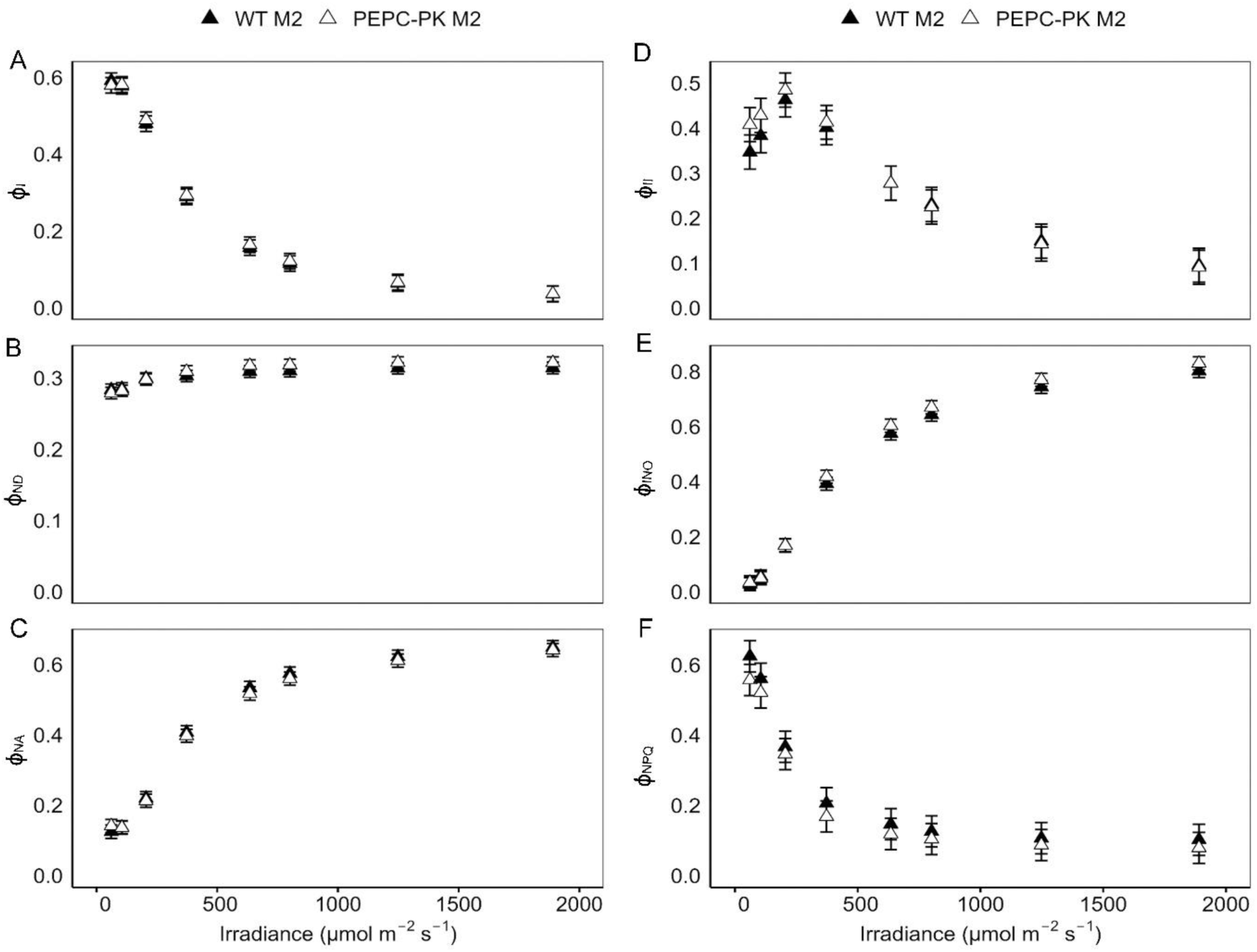
Light response curve of Photosystem I (PSI) and Photosystem II (PSII) related parameters in wild type and PEPC-PK mutant *Z. mays* genotype 2 (*ppck1-m2*). **(A-C)** The yield of photochemical reactions in PSI (ϕ_I_), non-photochemical loss due to the oxidized primary donor of PSI (ϕ_ND_), the non-photochemical yield of PSI due to acceptor side limitation (ϕ_NA_); **(D-F)** the effective quantum yield of PSII (ϕ_II_), quantum yield of non-regulated energy dissipation (ϕ_NO_), and the yield of non-photochemical quenching in PSII (ϕ_NPQ_). Mean ± SE, *n* = 6 biological replicates. No statistical differences were found between genotypes (Tukey-adjusted contrasts at *P* < 0.05).

**Supplementary Figure 5.**
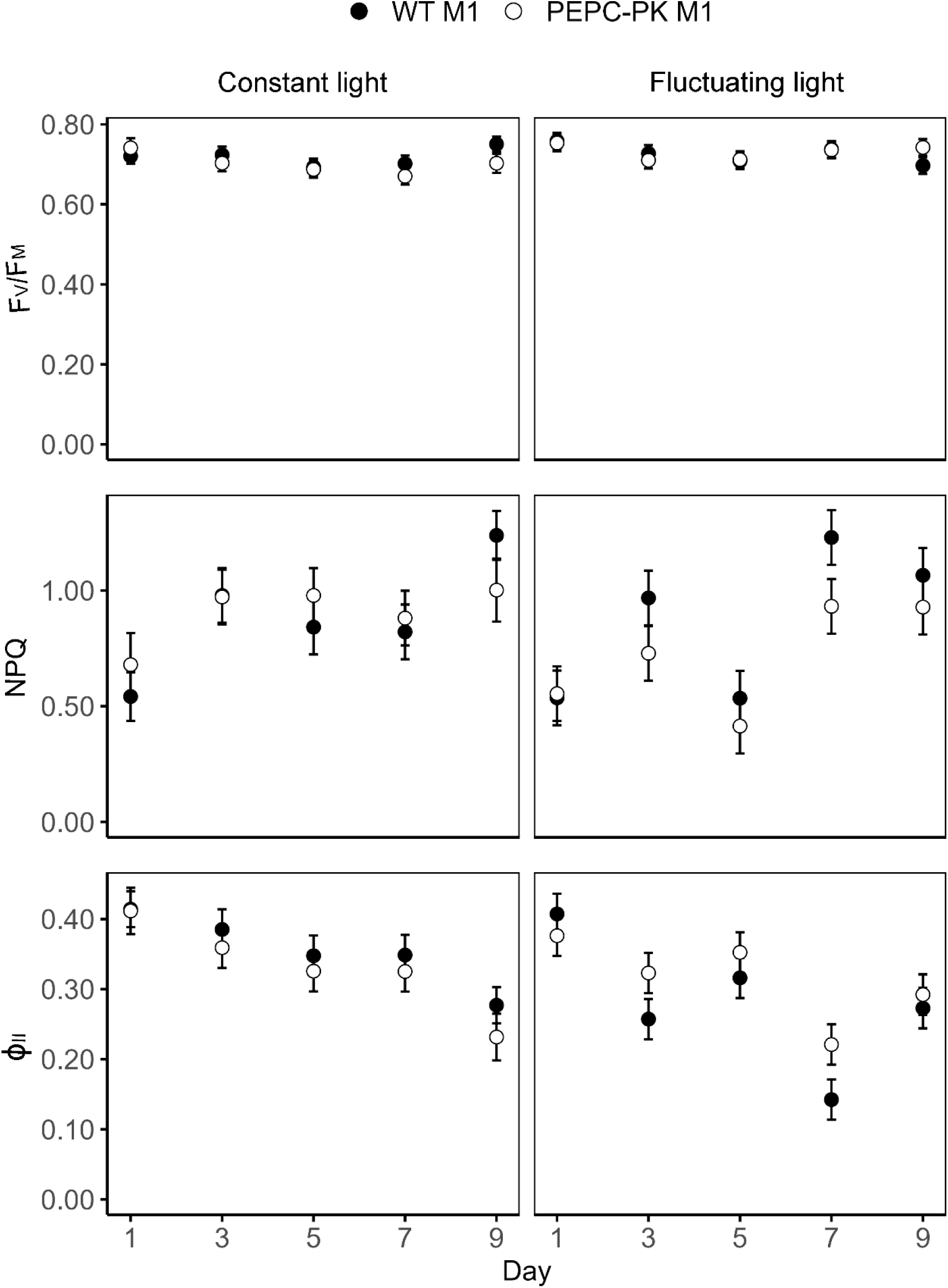
Effect of fluctuating light in WT M1 and PEPC-PK M1 (*ppck1-m1*) *Z. mays* plants on chlorophyll fluorescence parameters: maximum quantum efficiency of PSII (F_V_/F_M_), non-photochemical quenching (NPQ), and effective quantum yield of PSII (ϕ_II_) measured over 9 days under steady or fluctuating light conditions. Data represent means ± SE (n = 4). Different letters indicate significant differences between genotypes (Tukey-adjusted, *P* < 0.05).

**Supplementary Table 1.** ANOVA for the malate sensitivity of PEPC activity.

|  |  |  |  |  |
| --- | --- | --- | --- | --- |
| **Source of variation** | **df** | **Sum of squares** | **F value** | **Significance** |
| Genotype | 3 | 2190 | 17.6 | *** |
| Light condition | 1 | 4419 | 106.5 | *** |
| Malate concentration | 6 | 214050 | 859.8 | *** |
| Genotype × Light condition | 3 | 912 | 7.3 | ns |
| Genotype × Malate concentration | 18 | 979 | 1.3 | *** |
| Light condition × Malate concentration | 6 | 1648 | 6.6 | *** |
| Genotype × Light condition × Malate concentration | 18 | 282 | 0.3 | *** |
| Residuals | 161 | 6680 |  |  |

**Supplementary Table 2.** Analysis of variance (ANOVA) for PEPC activity in the absence of malate.

| **Source of variation** | **df** | **Sum of squares** | **F value** | **Significance** |
| --- | --- | --- | --- | --- |
| **Genotype** | 3 | 2321 | 0.9 | ns |
| **Light** | 1 | 7321 | 8.1 | ** |
| **Genotype × Light** | 3 | 751 | 0.3 | ns |
| **Residuals** | 24 | 21670 |  |  |

## References

Aldous SH, Weise SE, Sharkey TD, Waldera-Lupa DM, Stühler K, Mallmann J, Groth G, Gowik U, Westhoff P, Arsova B (2014) Evolution of the phosphoenolpyruvate bcarboxylase protein kinase family in C3 and C4 Flaveria spp. Plant Physiol. 165: 1076–1091

Arrivault S, Medeiros DB, Sales CRG, Guenther M, Kromdijk J, Fernie AR, Stitt M (2025) Metabolite profiling reveals slow and uncoordinated adjustment of C4 photosynthesis to sudden changes in irradiance. Plant Physiol. 199: kiaf508

Arrivault S, Obata T, Szecówka M, Mengin V, Guenther M, Hoehne M, Fernie AR, Stitt M (2017) Metabolite pools and carbon flow during C4 photosynthesis in maize: 13CO2 labeling kinetics and cell type fractionation. J. Exp. Bot. 68: 283–298

Baccolini C, Ishihara H, Feil R, Perez de Souza L, Alseekh S, Fernie AR, Stitt M, Lunn JE, Arrivault S (2025) Exploring the diversity of the CO_2_-concentrating mechanism (CCM) in different C4 subtypes. bioRxiv: 2025.2007.2015.664927

Bakrim N, Prioul J, Deleens E, Rocher J, Arrio-Dupont M, Vidal J, Gadal P, Chollet R (1993) Regulatory phosphorylation of C4 phosphoenolpyruvate carboxylase (a cardinal event influencing the photosynthesis rate in sorghum and maize). Plant Physiol. 101: 891–897

Bakrim Ni, Prioul J-L, Deleens E, Rocher J-P, Arrio-Dupont M, Vidal J, Gadal P, Chollet R (1993) Regulatory phosphorylation of C4 phosphoenolpyruvate carboxylase (a cardinal event influencing the photosynthesis rate in sorghum and maize). Plant Physiol. 101: 891–897

Balparda M, Bouzid M, Martinez MdP, Zheng K, Schwarzländer M, Maurino VG (2023) Regulation of plant carbon assimilation metabolism by post-translational modifications. The Plant Journal 114: 1059–1079

Bandarian V, Poehner WJ, Grover SD (1992) Metabolite activation of crassulacean acid metabolism and C4 phosphoenolpyruvate carboxylase. Plant Physiol. 100: 1411–1416

Bilger W, Björkman O (1990) Role of the xanthophyll cycle in photoprotection elucidated by measurements of light-induced absorbance changes, fluorescence and photosynthesis in leaves of *Hedera canariensis*. Photosynthesis Res. 25: 173–185

Boxall SF, Dever LV, Kneřová J, Gould PD, Hartwell J (2017) Phosphorylation of phospho enol pyruvate carboxylase is essential for maximal and sustained dark CO_2_ fixation and core circadian clock operation in the obligate crassulacean acid metabolism species *c*. The Plant Cell 29: 2519–2536

Britto DT, Kronzucker HJ (2005) Nitrogen acquisition, PEP carboxylase, and cellular pH homeostasis: new views on old paradigms. Plant, Cell Environ. 28: 1396–1409

Brutnell TP, Conrad LJ (2003) Transposon tagging using Activator (Ac) in maize. In Plant Functional Genomics. Springer, pp 157–175

Caburatan L, Park J (2021) Differential expression, tissue-specific distribution, and posttranslational controls of phosphoenolpyruvate carboxylase. Plants 10: 1887

Carter P, Nimmo H, Fewson C, Wilkins M (1990) *Bryophyllum fedtschenkoi* protein phosphatase type 2A can dephosphorylate phosphoenolpyruvate carboxylase. FEBS Lett. 263: 233–236

Carvalho P, Gomes C, Saibo NJ (2023) C4 Phosphoenolpyruvate Carboxylase: Evolution and transcriptional regulation. Genet. Mol. Biol. 46: e20230190

Chollet R, Vidal J, O’Leary MH (1996) Phosphoenolpyruvate carboxylase: a ubiquitous, highly regulated enzyme in plants. Annu. Rev. Plant Biol. 47: 273–298

Collatz GJ, Ribas-Carbo M, Berry JA (1992) Coupled photosynthesis-stomatal conductance model for leaves of C4 plants. Funct. Plant Biol. 19: 519–538

Cousins AB, Badger MR, Von Caemmerer S (2006) Carbonic anhydrase and its influence on carbon isotope discrimination during C4 photosynthesis. Insights from antisense RNA in *Flaveria bidentis*. Plant Physiol. 141: 232–242

Cousins AB, Baroli I, Badger MR, Ivakov A, Lea PJ, Leegood RC, Von Caemmerer S (2007) The role of phosphoenolpyruvate carboxylase during C4 photosynthetic isotope exchange and stomatal conductance. Plant Physiol. 145: 1006–1017

Cousins AB, Ghannoum O, Von Caemmerer S, Badger M (2010) Simultaneous determination of Rubisco carboxylase and oxygenase kinetic parameters in *Triticum aestivum* and *d* using membrane inlet mass spectrometry. Plant, Cell Environ. 33: 444–452

De la Osa C, Pérez-López J, Feria AB, Baena G, Marino D, Coleto I, Pérez-Montaño F, Gandullo J, Echevarría C, García-Mauriño S (2022) Knock-down of phosphoenolpyruvate carboxylase 3 negatively impacts growth, productivity, and respfonses to salt stress in sorghum (*Sorghum bicolor* L.). The Plant Journal 111: 231–249

Doncaster HD, Leegood RC (1987) Regulation of phosphoenolpyruvate carboxylase activity in maize leaves. Plant Physiol. 84: 82–87

Duff SM, Andreo CS, Pacquit V, Lepiniec L, Sarath G, Condon SA, Vidal J, Gadal P, Chollet R (1995) Kinetic analysis of the non-phosphorylated, in vitro phosphorylated, and phosphorylation-site-mutant (Asp8) forms of intact recombinant C4 phosphoenolpyruvate carboxylase from sorghum. Eur. J. Biochem. 228: 92–95

Ermakova M, Arrivault S, Giuliani R, Danila F, Alonso-Cantabrana H, Vlad D, Ishihara H, Feil R, Guenther M, Borghi GL (2021) Installation of C4 photosynthetic pathway enzymes in rice using a single construct. Plant Biotechnol. J. 19: 575–588

Ermakova M, Danila FR, Furbank RT, von Caemmerer S (2020) On the road to C4 rice: advances and perspectives. The Plant Journal 101: 940–950

Fryer MJ, Andrews JR, Oxborough K, Blowers DA, Baker NR (1998) Relationship between CO_2_ assimilation, photosynthetic electron transport, and active O_2_ metabolism in leaves of maize in the field during periods of low temperature. Plant Physiol. 116: 571–580

Fukayama H, Hatch MD, Tamai T, Tsuchida H, Sudoh S, Furbank RT, Miyao M (2003) Activity regulation and physiological impacts of maize C4-specific phospho enol pyruvate carboxylase overproduced in transgenic rice plants. Photosynthesis Res. 77: 227–239

Fukayama H, Tamai T, Taniguchi Y, Sullivan S, Miyao M, Nimmo HG (2006) Characterization and functional analysis of phosphoenolpyruvate carboxylase kinase genes in rice. The plant journal 47: 258–268

Furumoto T, Izui K, Quinn V, Furbank RT, von Caemmerer S (2007) Phosphorylation of phospho enol pyruvate carboxylase is not essential for high photosynthetic rates in the C4 species Flaveria bidentis. Plant Physiol. 144: 1936–1945

Furumoto T, Izui K, Quinn V, Furbank RT, Von Caemmerer S (2007) Phosphorylation of phosphoenolpyruvate carboxylase is not essential for high photosynthetic rates in the C4 species *Flaveria bidentis*. Plant Physiol. 144: 1936–1945

Gao Y, Woo K (1996) Regulation of phosphoenolpyruvate carboxylase in *Zea mays* by protein phosphorylation and metabolites and their roles in photosynthesis. Aust. J. Plant Physiol. 23: 25–32

Genty B, Briantais J-M, Baker NR (1989) The relationship between the quantum yield of photosynthetic electron transport and quenching of chlorophyll fluorescence. Biochim Biophys Actai 990: 87–92

Ghannoum O (2009) C4 photosynthesis and water stress. Ann. Bot. 103: 635–644

Giuliani R, Karki S, Covshoff S, Lin H-C, Coe RA, Koteyeva NK, Evans MA, Quick WP, von Caemmerer S, Furbank RT (2019) Transgenic maize phosphoenolpyruvate carboxylase alters leaf–atmosphere CO_2_ and ^13^CO_2_ exchanges in *Oryza sativa*. Photosynthesis Res. 142: 153–167

Grieco M, Tikkanen M, Paakkarinen V, Kangasjärvi S, Aro E-M (2012) Steady-state phosphorylation of light-harvesting complex II proteins preserves photosystem I under fluctuating white light. Plant Physiol. 160: 1896–1910

Hartwell J, Jenkins G, Wilkins M, Nimmo H (1999) The light induction of maize phosphoenolpyruvate carboxylase kinase translatable mRNA requires transcription but not translation. Plant, Cell Environ. 22: 883–889

Hatch MD (1987) C4 photosynthesis: a unique elend of modified biochemistry, anatomy and ultrastructure. Biochimica et Biophysica Acta (BBA)-Reviews on Bioenergetics 895: 81–106

Hibberd JM, Covshoff S (2010) The regulation of gene expression required for C4 photosynthesis. Annu. Rev. Plant Biol. 61: 181–207

Huber SC, Sugiyama T (1986) Changes in sensitivity to effectors of maize leaf phosphoenolypyruvate carboxylase during light/dark transitions. Plant Physiol. 81: 674–677

Izui K, Matsumura H, Furumoto T, Kai Y (2004) Phospho enol pyruvate carboxylase: a new era of structural biology. Annu. Rev. Plant Biol. 55: 69–84

Jiao J-A, Chollet R (1989) Regulatory seryl-phosphorylation of C4 phosphoenolpyruvate carboxylase by a soluble protein kinase from maize leaves. Archives of biochemistry and biophysics 269: 526–535

Jiao J-a, Chollet R (1990) Regulatory phosphorylation of serine-15 in maize phosphoenolpyruvate carboxylase by a C4-leaf protein-serine kinase. Archives of biochemistry and biophysics 283: 300–305

Jiao J-a, Echevarría C, Vidal J, Chollet R (1991) Protein turnover as a component in the light/dark regulation of phosphoenolpyruvate carboxylase protein-serine kinase activity in C4 plants. Proceedings of the National Academy of Sciences 88: 2712–2715

Kajala K, Covshoff S, Karki S, Woodfield H, Tolley BJ, Dionora MJA, Mogul RT, Mabilangan AE, Danila FR, Hibberd JM (2011) Strategies for engineering a two-celled C4 photosynthetic pathway into rice. J. Exp. Bot. 62: 3001–3010

Klughammer C, Schreiber U (2008) Complementary PS II quantum yields calculated from simple fluorescence parameters measured by PAM fluorometry and the Saturation Pulse method. PAM application notes 1: 201–247

Kramer DM, Johnson G, Kiirats O, Edwards GE (2004) New fluorescence parameters for the determination of QA redox state and excitation energy fluxes. Photosynthesis Res. 79: 209–218

Lenth R (2023) emmeans: Estimated Marginal Means, aka Least-Squares Means. R package version 1.8. 5

Lin H, Arrivault S, Coe RA, Karki S, Covshoff S, Bagunu E, Lunn JE, Stitt M, Furbank RT, Hibberd JM (2020) A partial C4 photosynthetic biochemical pathway in rice. Front. Plant Sci. 11: 564463

Meimoun P, Gousset-Dupont A, Lebouteiller B, Ambard-Bretteville F, Besin E, Lelarge C, Mauve C, Hodges M, Vidal J (2009) The impact of PEPC phosphorylation on growth and development of Arabidopsis thaliana: molecular and physiological characterization of PEPC kinase mutants. FEBS Lett. 583: 1649–1652

Muñoz-Clares RA, González-Segura L, Juárez-Díaz JA, Mújica-Jiménez C (2020) Structural and biochemical evidence of the glucose 6-phosphate-allosteric site of maize C4-phosphoenolpyruvate carboxylase: its importance in the overall enzyme kinetics. Biochem. J. 477: 2095–2114

Nimmo HG (2000) The regulation of phosphoenolpyruvate carboxylase in CAM plants. Trends Plant Sci. 5: 75–80

Nimmo HG (2003) Control of the phosphorylation of phosphoenolpyruvate carboxylase in higher plants. Archives of Biochemistry and Biophysics 414: 189–196

Nimmo HG, Fontaine V, Hartwell J, Jenkins GI, Nimmo GA, Wilkins MB (2001) PEP carboxylase kinase is a novel protein kinase controlled at the level of expression. New Phytol. 151: 91–97

Nishikido T, Takanashi H (1973) Glycine activation of PEP carboxylase from monocotyledoneous C4 plants. Biochem. Biophys. Res. Commun. 53: 126–133

O’Leary B, Park J, Plaxton WC (2011) The remarkable diversity of plant PEPC (phosphoenolpyruvate carboxylase): recent insights into the physiological functions and post-translational controls of non-photosynthetic PEPCs. Biochem. J. 436: 15–34

Peisker M (1979) Conditions for low, and oxygen-independent, CO_p_ compensation concentrations in C4 plants as derived from a simple model.

Pengelly JJ, Sirault XR, Tazoe Y, Evans JR, Furbank RT, Von Caemmerer S (2010) Growth of the C4 dicot *Flaveria bidentis*: photosynthetic acclimation to low light through shifts in leaf anatomy and biochemistry. J. Exp. Bot. 61: 4109–4122

Pinheiro J (2011) nlme: Linear and nonlinear mixed effects models. R package version 3: 1

Ritz C, Baty F, Streibig JC, Gerhard D (2015) Dose-response analysis using R. PloS one 10: e0146021

Sage RF (2004) The evolution of C4 photosynthesis. New Phytol. 161: 341–370

Sage RF, Sage TL, Kocacinar F (2012) Photorespiration and the evolution of C4 photosynthesis. Annu. Rev. Plant Biol. 63: 19–47

Sánchez R, Cejudo FJ (2003) Identification and expression analysis of a gene encoding a bacterial-type phosphoenolvpyruvate carboxylase from Arabidopsis and rice. Plant Physiol. 132: 949–957

Slattery RA, Walker BJ, Weber AP, Ort DR (2018) The impacts of fluctuating light on crop performance. Plant Physiol. 176: 990–1003

Studer AJ, Gandin A, Kolbe AR, Wang L, Cousins AB, Brutnell TP (2014) A limited role for carbonic anhydrase in C4 photosynthesis as revealed by a ca1ca2 double mutant in maize. Plant Physiol. 165: 608–617

Sun W, Ubierna N, Ma J-Y, Walker BJ, Kramer DM, Cousins AB (2014) The coordination of C4 photosynthesis and the CO2-concentrating mechanism in maize and Miscanthus× giganteus in response to transient changes in light quality. Plant Physiol. 164: 1283–1292

Svensson P, Bläsing OE, Westhoff P (2003) Evolution of C4 phosphoenolpyruvate carboxylase. Archives of Biochemistry and Biophysics 414: 180–188

Taniguchi Y, Ohkawa H, Masumoto C, Fukuda T, Tamai T, Lee K, Sudoh S, Tsuchida H, Sasaki H, Fukayama H (2008) Overproduction of C4 photosynthetic enzymes in transgenic rice plants: an approach to introduce the C4-like photosynthetic pathway into rice. J. Exp. Bot. 59: 1799–1809

Terada K, Kai T, Okuno S, Fujisawa H, Izui K (1990) Maize leaf phosphoenolpyruvate carboxylase: phosphorylation of Ser15 with a mammalian cyclic AMP-dependent protein kinase diminishes sensitivity to inhibition by malate. FEBS Lett. 259: 241–244

Tovar-Méndez A, Mujica-Jiménez C, Munoz-Clares RA (2000) Physiological implications of the kinetics of maize leaf phospho enol pyruvate carboxylase. Plant Physiol. 123: 149–160

Ueno Y, Imanari E, Emura J, Yoshizawa-Kumagaye K, Nakajima K, Inami K, Shiba T, Sakakibara H, Sugiyama T, Izui K (2000) Immunological analysis of the phosphorylation state of maize C4-form phosphoenolpyruvate carboxylase with specific antibodies raised against a synthetic phosphorylated peptide. The Plant Journal 21: 17–26

Vidal J, Chollet R (1997) Regulatory phosphorylation of C4 PEP carboxylase. Trends Plant Sci. 2: 230–237

von Caemmerer S (2000) Biochemical models of leaf photosynthesis. Csiro publishing

von Caemmerer S (2021) Updating the steady-state model of C4 photosynthesis. J. Exp. Bot. 72: 6003–6017

von Caemmerer S, Furbank RT (1999) Modeling C4 photosynthesis. C4 plant biology: 173–211

von Caemmerer S, Furbank RT (2003) The C4 pathway: an efficient CO_2_ pump. Photosynthesis Res. 77: 191–207

von Caemmerer S, Quick WP, Furbank RT (2012) The development of C4 rice: current progress and future challenges. science 336: 1671–1672

Walker BJ, VanLoocke A, Bernacchi CJ, Ort DR (2016) The costs of photorespiration to food production now and in the future. Annu. Rev. Plant Biol. 67: 107–129

Weber AP, von Caemmerer S (2010) Plastid transport and metabolism of C3 and C4 plants—comparative analysis and possible biotechnological exploitation. Curr. Opin. Plant Biol. 13: 256–264

Wickham H (2016) ggplot2: Elegant Graphics for Data Analysis. New York: SpringerVerlag. Retrieved May 28: 2022

